# TonB-dependent Transporters of Fe-petrobactin in *Marinobacter nauticus* and Southern Ocean bacteria

**DOI:** 10.64898/2026.10.08.756784

**Authors:** Sophie Nolivos, Rhea Thoppil, Florence Hakil, Audrey Guéneuguès, Philippe Catala, Katarzyna Kinksa, Ryszard Lobinski, Stéphane Blain, Ingrid Obernosterer, Régis Grimaud

## Abstract

TonB-dependent transporters (TBDTs) play a central role in siderophore-mediated iron acquisition in the marine environment. However, assigning substrate specificity to TBDTs remains a major bottleneck in understanding the ecological and mechanistic roles of siderophore production and uptake. Here, we identify in *Marinobacter nauticus* two TBDTs, OptA and OptB, that share low sequence identity yet both mediate Fe-petrobactin uptake. Despite this functional overlap, they underlie distinct regulatory mechanisms, suggesting differential utilization under specific environmental conditions. Phylogenetic analyses indicate that OptA likely co-evolved with the petrobactin biosynthetic gene cluster, whereas OptB appears to have followed a distinct evolutionary trajectory, acquiring the capacity to import petrobactin as a xenosiderophore.

Using resuscitated communities from the Southern Ocean, we further show that Fe-petrobactin stimulated prokaryotic growth. Based on amino acid similarity, we identified OptB-like transporters encoded in metagenomes and MAGs and show that taxa harbouring these transporters, such as *Pseudoaltermonas*, respond specifically to Fe-petrobactin addition. OptB-like transporters were also detected in environmental metagenomes from the Southern Ocean, whereas the petrobactin biosynthetic gene clusters were not, indicating that petrobactin primarily functions as a xenosiderophore in natural communities.

## Introduction

Iron is a vital micronutrient for bacterial growth because it acts as a cofactor in enzymes involved in key metabolic pathways such as respiration, DNA synthesis, and the detoxification of reactive oxygen species. However, in many environments, iron acquisition is challenging due to the low ambient concentrations, chemical forms that cannot be directly assimilated and competition with other microorganisms. Together, these factors create vast iron-limited regions, such as the Southern Ocean (SO), where iron bioavailability profoundly affects microbial community composition, activity, and element cycling [1,2].

To overcome iron scarcity, marine bacteria have evolved diverse strategies, with siderophore-mediated uptake playing a central role. Siderophores are low-molecular-weight organic ligands, that chelate Fe³⁺ with exceptionally high stability constants (log K ≈ 30–50) [3]. Once secreted, siderophores bind and solubilize Fe^3+^ or displace it from other ligands enabling uptake under limiting conditions. In Gram-negative bacteria, Fe-siderophore complex uptake is mediated by TonB-dependent transporters (TBDTs) [4,5].

Marine siderophores exhibit remarkable structural and functional diversity, with over 80 families identified, including ferrioxamines, amphibactins, synechobactins, and petrobactin, among the most frequently detected in seawater [6–9]. Metagenomic and metatranscriptomic surveys show that genes encoding siderophore biosynthesis and uptake are widely distributed throughout the ocean [9–15], with TBDT genes generally more abundant than siderophore biosynthesis genes. This suggests that marine bacteria can exploit multiple siderophores, including xenosiderophores (produced by other organisms), provided they express the appropriate TBDTs [3,13–16]. Consequently, siderophores represent public goods that shape community structure through cooperative, exploitative, and competitive interactions. While such interactions are well documented under laboratory conditions, evidence in marine environments remains limited [16,17].

The ecological role of siderophores depends on the populations that can utilize them through the expression of specific TBDTs. However, TBDTs are highly divergent at the sequence level, complicating the prediction of substrate specificity. Few marine siderophore transporters have been experimentally characterized, primarily in *Vibrio* species [18], leaving most TBDT genes in marine bacterial genomes and metagenomes unassigned to specific siderophores. This study focuses on Fe-petrobactin transport in *Marinobacter nauticus* and SO bacterial communities. Petrobactin is a mixed-type siderophore comprising catecholate and α-hydroxycarboxylate functional groups, originally characterized in *Marinobacter nauticus* [19]. Petrobactin increases iron bioavailability and is widely distributed in marine environments [8–10]. The biosynthetic pathway was elucidated in *Bacillus anthracis* [20] and described in *Alteromonas macleodii* and *M. nauticus* [8,21]. In the present study, we identified two transporters in *M. nauticus*, OptA and OptB. Genomic context and expression analyses indicate that OptA is associated with petrobactin biosynthesis whereas *optB*-like genes were identified in SO communities that utilise Fe-petrobactin but lack a detectable petrobactin biosynthetic gene cluster, suggesting a function for xenosiderophore uptake.

## Materials and Methods

### Bacterial strains, plasmids, chemicals and growth conditions

Bacterial strains, plasmids and primers used in this study are listed in **Tables S1**, **S2** and **S3**, respectively. *M. nauticus* JM1 strain was used as the wild-type strain [22]. Gene deletion was previously described [21]. *M. nauticus* strains were routinely cultivated in Halo medium or EMS medium [22]. All experiments, requiring a controlled iron concentration, were performed in iron-free MEM medium as previously described [21]. All chemicals were from Sigma-Aldrich, unless otherwise specified.

### Supplementation of TBDTs mutants with petrobactin

*M. nauticus* strains were streaked from –80°C glycerol stock onto fresh EMS (7.2 µM FeCl_2_) plates containing 20 mM lactate and 100 μM streptomycin and incubated at 30°C for five days. A few colonies were then resuspended in iron-free MEM and used to inoculate cultures at an A_600nm_ of 0.005. The cultures were performed in MEM medium supplemented with 1 µM FeCl₃, 200 µM EDTA and when required, 10 µM petrobactin (Supplementary Information) which was prepared one day in advance. At the time of inoculation, 20 mM of acetate were added to the cultures, which were then incubated at 30°C and 200 rpm.

### RNA extraction and Reverse Transcription quantitative PCR (RT-qPCR)

RNA extraction and RT-qPCR were performed as previously described [21]. Briefly, *M. nauticus* was cultured in triplicate in MEM acetate with 7 200 or 200 nmol L⁻¹ FeCl_3_ until A_600nm_ reached 0.4 or after 45 h of growth. Two milliliters of cells were harvested and resuspended in 200 µL of 10 mmol L⁻¹ Tris-HCl before RNAs extraction. For qPCR reactions, 1.25 µg of cDNA synthesized from 325 ng of RNA was used. See Supplementary Information for details.

### Phylogenetic analysis of TBDT

Sixty-five TBDT sequences encoded by genes located in the genomic vicinity of predicted petrobactin-related biosynthetic gene clusters were retrieved (see Supplementary Information). Sequences belonging to the same genus or species and sharing more than 90% identity were represented by a single sequence. The best-conserved homolog of OptA from *Alteromonas macleodii* ATCC 27126 (MASE_17930) together with a subset of TBDT sequences with experimentally determined siderophore substrates retrieved from the UniProt database [23] and the RCSB Protein Data Bank [24] were added to the data set. Then, signal peptide sequences were removed based on cleavage site predictions obtained with SignalP 6.0 [25]. The resulting mature protein sequences were aligned using MUSCLE. This alignment was used to construct a phylogenetic tree using the Neighbor-Joining method [26] with the JTT matrix-based model [27], implemented in MEGA 12 [28].

### Sampling and cryopreservation of Southern Ocean microbial communities

Seawater was collected at two stations during the SWINGS cruise (GEOTRACES GS02; doi 10.17600/18001925) in the Indian Sector of the SO (**Fig. S1**). Seawater was collected from the ship’s hull intake, through the moonpool with a polytetrafluoroethylene (PTFE) diaphragm pump and tubing, and filtered through 0.8 µm polycarbonate (PC) filters (Nucleopore). Cells present in the < 0.8 µm size fraction were concentrated on cellulose nitrate filters (0.2 µm pore size, 47 mm diameter; Millipore)(5.45 L, Station 27) and on polyethersulfone filters (0.45 µm pore size, 25 mm diameter, Millipore)(2.55 L, Station 47) following the protocol of Rain-Franco et al. [29]. Each cell concentrate, corresponding to 2 mL, was then split into 2 subsamples, one for flow cytometry analyses and one for resuscitation. For flow cytometry, 800 µL were fixed with 1% glutaraldehyde, stored at 4 °C for 30 min before being shock-frozen in liquid nitrogen, and stored at –80 °C until analysis in the home lab. The remaining volumes were cryopreserved in 5% DMSO, stored at 4 °C for 15 min, shock-frozen in liquid nitrogen, and stored at –80°C until use. In addition, 20 L of surface seawater were collected in the Antarctic Zone using the GeoFISH [30] following trace metal clean procedures [31]. Seawater was 0.2 µm filtered in the trace metal clean container and transported in acid-cleaned PC bottles.

### Incubation experiments of resuscitated microbial communities

The cell concentrates were diluted with the collected seawater to obtain a cell abundance equal to 10% of the *in situ* abundance in surface waters. Experiments were performed in triplicates, with 100 ml of cultures amended with 10 µM trace metal free glucose, that was passed over a Chelex 100 ion-exchange resin (Bio-Rad, 200400 mesh). When indicated, Fe-petrobactin complexes were added at 10 nM (Station 27) or 6 nM (Station 47). Petrobactin was purified as describe previously [32]. Fe-petrobactin complexes preparation is described in the Supplementary Information. All culture work and subsampling were conducted in a clean room (Class 10,000) equipped with a laminar flow hood (Class 100). PC bottles and plasticware were soaked in 10% HCl for 24 h, and subsequently soaked overnight with ultrapure water. Incubations were carried out in the dark for 14 days at 4 °C. Subsamples for flow cytometry were taken every 24 h. Flow cytometry analyses were done on a FACSCanto II flow cytometer (Becton Dickinson) as described previously [33].

### Microbial community composition and metagenomics

At the end of the incubation, cells were concentrated on 0.2 µm PC filters (47 mm diameter, Nuclepore) and DNA extracted using the AllPrep Kit (Qiagen, Hilden, Germany) as described previously [34]. PCR amplification was performed using the primers 515F–Y (5′-GTGYCAGCMGCCGCGG TAA) and 926R (5′-CCGYCAATTYMTTTRAGTTT) [35]. The *16S rDNA* gene amplicons were sequenced via next-generation sequencing (Illumina MiSeq 2 × 250 bp chemistry on one flow cell) on the LGBT platform (Berlin, Germany). A Mock community DNA (LGC standards, UK) was used as a standard for subsequent analyses and was considered as a DNA sample for all treatments. Raw sequences were demultiplexed and quality filtered using the Illumina bcl2fastq v.2.20 at the platform Biosearch Technologies (Berlin, Germany). The total number of reads per sample are provided in **Table S4**. To achieve random subsampling, samples were rarefied to 20 212 reads in the PFZ experiment and to 28 534 reads in the AAZ experiment using *phyloseq* v1.40 [36] in *R*. Taxonomy was assigned to the amplicon sequence variants (ASVs) based on the GTDB R220 reference database [37] using *DADA2* v1.24 [38] in *R* v4.2.1 [39]. A total of 120 ASVs were obtained for both experiments (**Table S5**); only ASVs with a relative abundance of > 1% in at least one sample were considered for further analyses.

Metagenomic samples were sequenced using an Illumina NovaSeq 6000 system with 2 × 150 bp chemistry at Fasteris SA, Inc. (Switzerland). Each sample yielded 28-40 million pairs of metagenomic reads, totalling 261.8 GB (**Table S6**). Raw reads were checked using FastQC and processed using *trimmomatic* v0.32 [40]. Because the microbial communities for the inoculation of the two experiments were sampled at different locations (**Fig. S1**), a co-assembly metagenome approach was carried out for each of the experiments. The bioinformatics pipeline is detailed in the Supplementary Information.

#### Statistical analyses and visualization of data

Differences in growth curves between the control and Fe-petrobactin amended incubations were tested using a linear mixed-effects model (LMM) using *lme4* v1.1.37 [41] and model significance was assessed using analysis of variance (ANOVA; [42]). Visualization of all the above-mentioned data was performed using *ggplot2* v3.5.1 and heatmaps were produced using *pheatmap* v1.0.12 in R, using the Euclidean distance measure for distance (v4.1.2; [39]). Differences in microbial community composition were determined using analysis of similarities (ANOSIM; [43]) based on Bray–Curtis dissimilarity [44] using *phyloseq* v1.40 [36] and *vegan* v2.6 [45].

## Results

### Two TBDTs transport the petrobactin in *M. nauticus*

The *M. nauticus* genome encodes six proteins containing the Pfam00593 domain, characteristic of TBDTs. A BLAST search for these six putative TBDTs against the UniProtKB/Swiss-Prot database revealed low sequence conservation (19.8–25.4% identity) relative to TBDTs with experimentally characterised substrates (**Table S7**). This finding highlights the substantial sequence divergence among TBDTs, which complicates the reliable prediction of substrate specificity.

Examination of the genomic context of the six TBDT-encoding genes provided insights into their potential functions. Three putative iron-related TBDTs, hereafter designated OptA (Outer membrane Petrobactin Transporter A), OptB, and FhuE, as well as one vitamin B12 transporter (BtuB), were identified. The *optA*, *optB*, and *fhuE* genes are located adjacent to genes encoding a putative Fe-siderophore ABC transporter. Notably, *optA* lies in close proximity to the *asbABCDEF* petrobactin biosynthesis cluster (**Fig. 1**).

**Figure 1.**
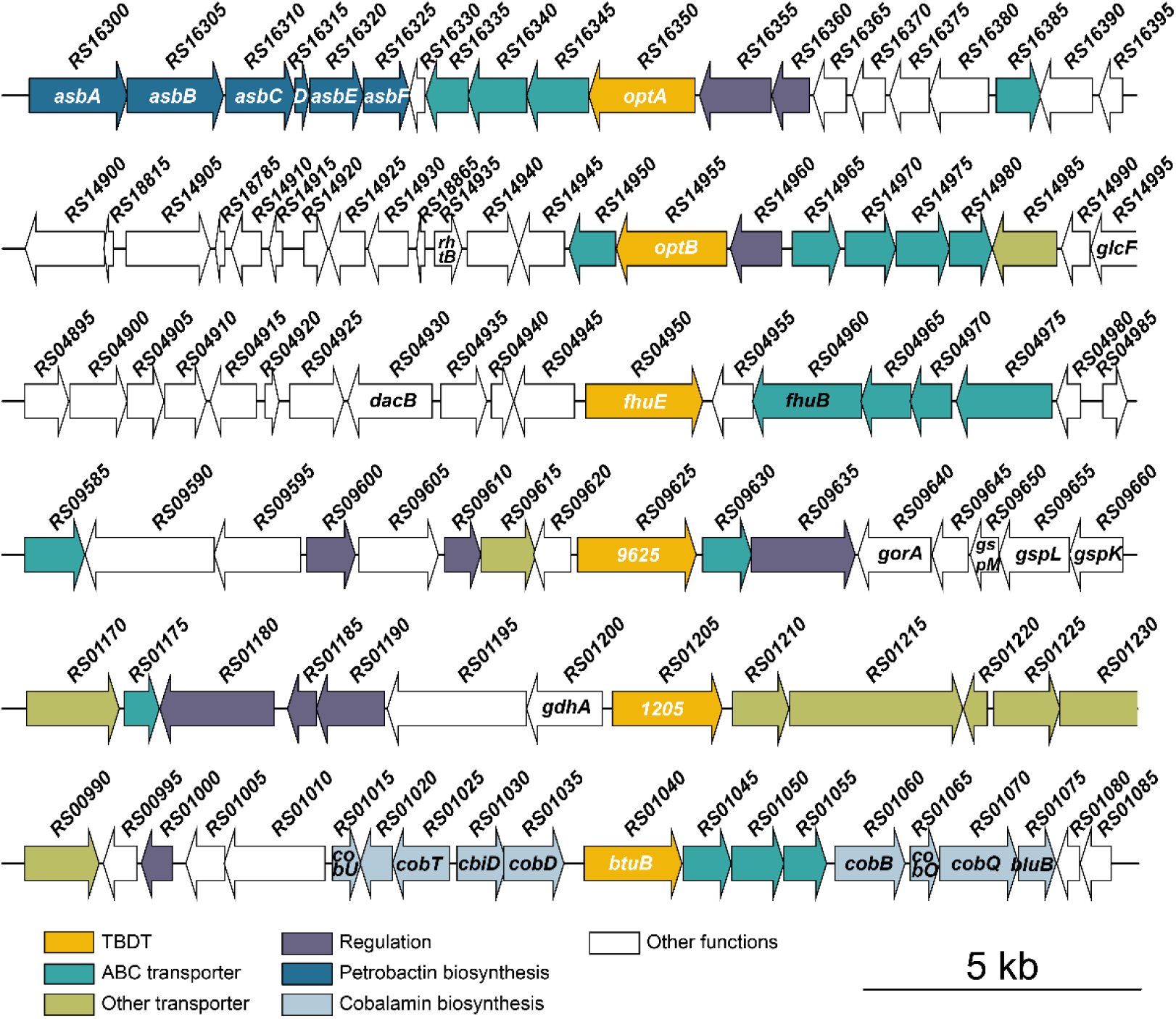
G**e**nomic **context of the six putative TBDT genes in *M. nauticus* SP17.** Arrows indicate gene length and orientation along the chromosome. Colours denote predicted functions. Gene names shown in black correspond to GenBank annotations, whereas those in white were assigned in this study based on BLAST analyses, global sequence alignments (**Tables S7 and S8**), and the results presented here. Full locus tags in the GeneBank Reference Sequence NC_017067.1 begin with ‘MARHY_’.

Genes involved in iron acquisition are typically induced under low-iron conditions. To further elucidate the role of TBDTs in iron acquisition, we compared the expression levels of TBDT-encoding genes at high and low-iron concentrations (**Fig. 2**). Among the six candidate genes, three (*optA optB* and *fhuE)* were overexpressed under low-iron conditions, suggesting their involvement in iron acquisition.

**Figure 2.**
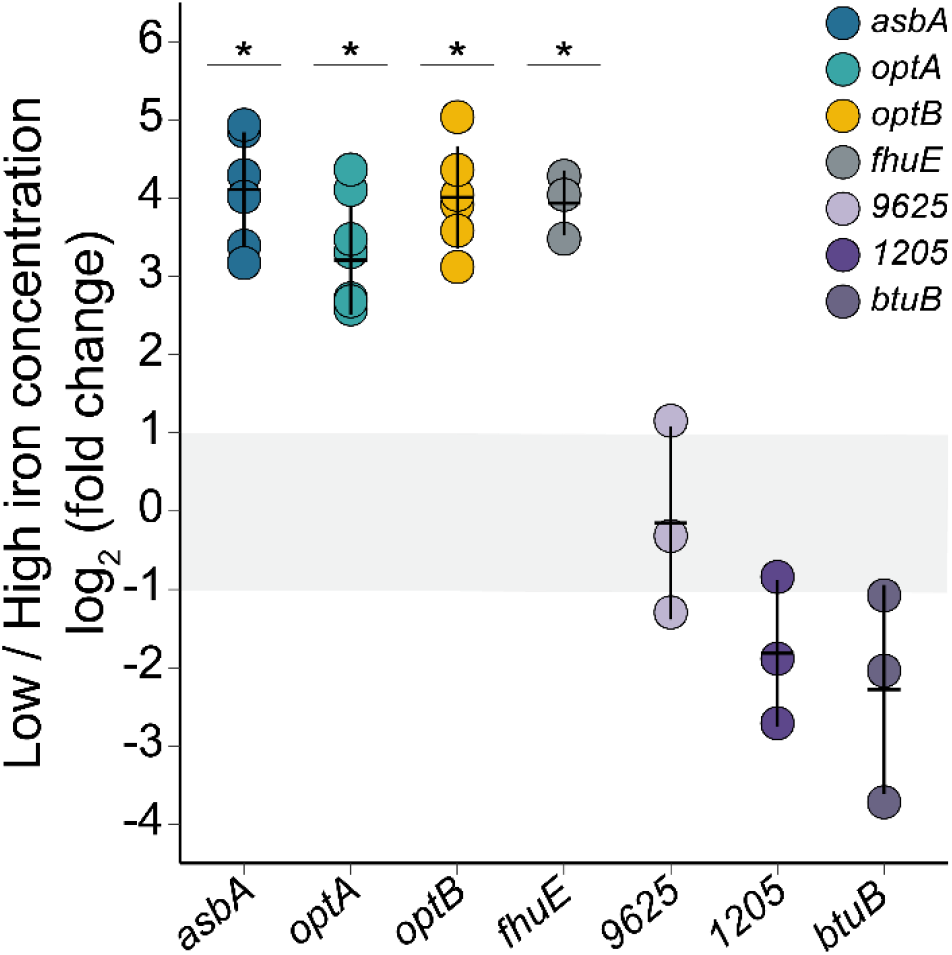
E**x**pression **ratios of *asbA* and the six TBDT-encoding genes under low versus high iron concentration.** log_2_ ratios of the expression levels measured by RT-qPCR at 200 nmol·L⁻¹ of iron relative to 7 200 nmol·L⁻¹ of iron and normalised to *rpoD* gene expression. Relative fold changes in gene expression were obtained according to the 2^-ΔΔCt^ method. Each experiment was performed in at least triplicate. Each dot represents the mean of three RT-qPCR reactions performed on one biological replicate. Solid lines represent the mean ± standard deviation (SD) of biological replicates. Asterisks indicate log_2_ fold change values significantly different from 1, absence of asterisk indicates log_2_ fold change values not different from 1, 0 or –1 (*p* < 0.05, one-sample t-test).

To determine which TBDT mediates petrobactin uptake, we assessed the ability of TBDT mutants to utilise petrobactin. First, a mutant lacking the petrobactin biosynthesis genes (Δ*asbABF*) failed to grow when iron was supplied as an Fe-EDTA chelate (**Fig. 3**). However, petrobactin supplementation restored growth, indicating that under these conditions *M. nauticus* relies on petrobactin for iron acquisition and therefore possesses the transporter (**Fig. 3**). All mutants deficient in the petrobactin biosynthesis genes and one of the iron-related TBDT genes (*optA*, *opB* and *fhuE*) were still rescued by petrobactin addition (**Fig. 3, Fig. S2**). It was concluded that none of these TBDTs function as a petrobactin transporter, or that petrobactin is transported by multiple TBDTs. To test the latter hypothesis, deletions of two TBDT genes were constructed in the Δ*asbABF* background. The growth of the Δ*asbABF* Δ*optA* Δ*optB* mutant was not restored by the addition of petrobactin **(Fig. 3)**, demonstrating that both OptA and OptB are required for petrobactin utilisation. In contrast, the Δ*asbABF* Δ*optA* Δ*fhuE* mutant was rescued, indicating that FhuE is not involved in petrobactin transport. The involvement of TBDTs 1205 and 9625, as well as BtuB, was likewise excluded (**Fig. S2**).

**Figure 3.**
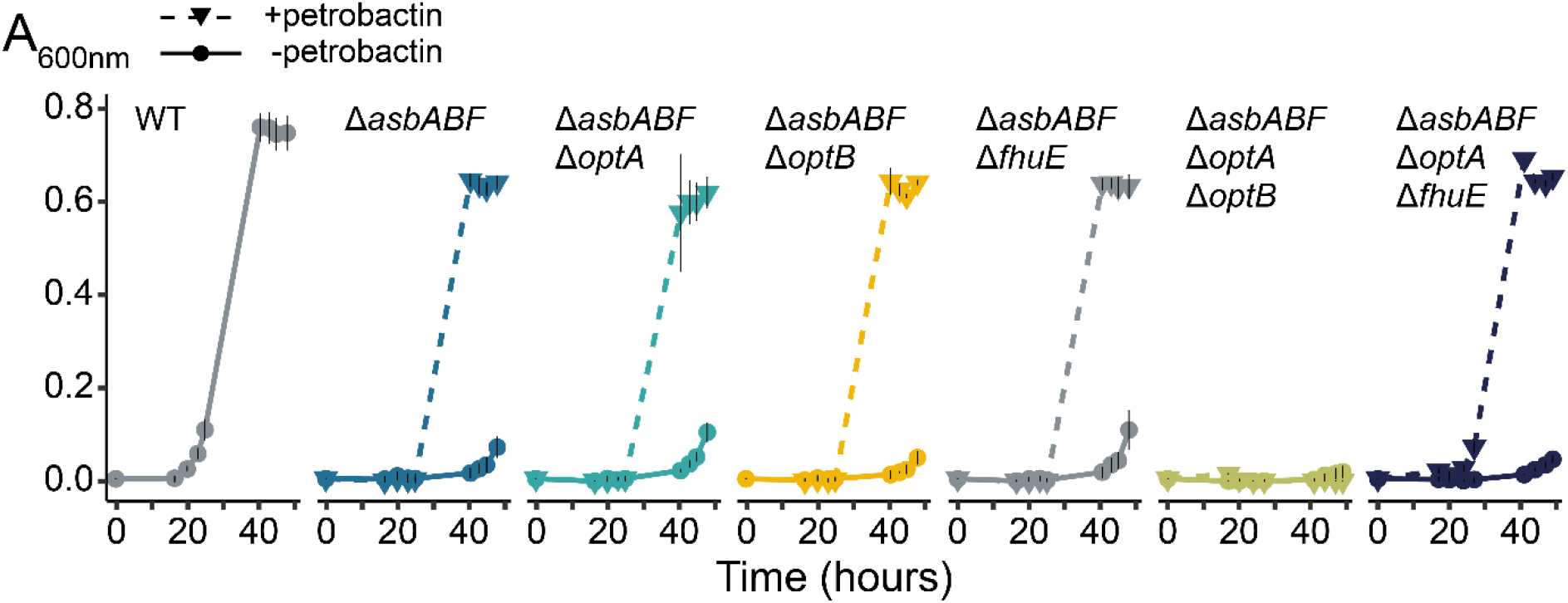
P**e**trobactin **utilisation by TBDT deletion mutants.** Growth curves of the wild type (WT), a petrobactin-production-deficient mutant (Δ*asbABF)* and single– or double-TBDT deletion mutant derivative, in the presence (dashed lines) or absence (solid lines) of 10 µM purified petrobactin. WT (JM1), Δ*asbABF* (FH35), Δ*asbABF* Δ*optA* (FH46), Δ*asbABF* Δ*optB* (FH50), Δ*asbABF* Δ*fhuE* (FH51), Δ*asbABF* Δ*optA* Δ*optB* (FH71), Δ*asbABF* Δ*optA* Δ*fhuE* (FH72).

These supplementation experiments demonstrate that petrobactin is exclusively transported by OptA and OptB.

### Differential cellular control of OptA and OptB production

It is well established that the expression of some TBDTs is regulated by the presence of their cognate iron-loaded siderophore [46]. We therefore examined the expression of the three iron-responsive TBDT genes (*optA*, *optB*, and *fhuE*) in a *M. nauticus* strain deficient in petrobactin biosynthesis (Δ*asbABF*), both in the presence or absence of petrobactin. Petrobactin addition failed to induce the expression of any of these transporter genes (**Fig. S3A**). These results suggest that TBDT expression is either independent of petrobactin availability or that alternative regulatory mechanisms compensate for the lack of the siderophore in the mutant strain. To further investigate the regulation of iron uptake, we analysed gene induction under low-iron conditions in mutants lacking iron transport functions (**Fig. 4A**). Deletion of either the petrobactin biosynthesis genes (Δ*asbABF*) or the transporter gene *optA* abolished the induction of other iron uptake genes under iron starvation **(Fig. 4A**), indicating that both petrobactin production and the presence of OptA are required for iron-responsive regulation. In contrast, OptB and FhuE did not contribute to this regulatory response, indicating distinct roles for OptA and OptB in the cellular response to iron limitation.

**Figure 4.**
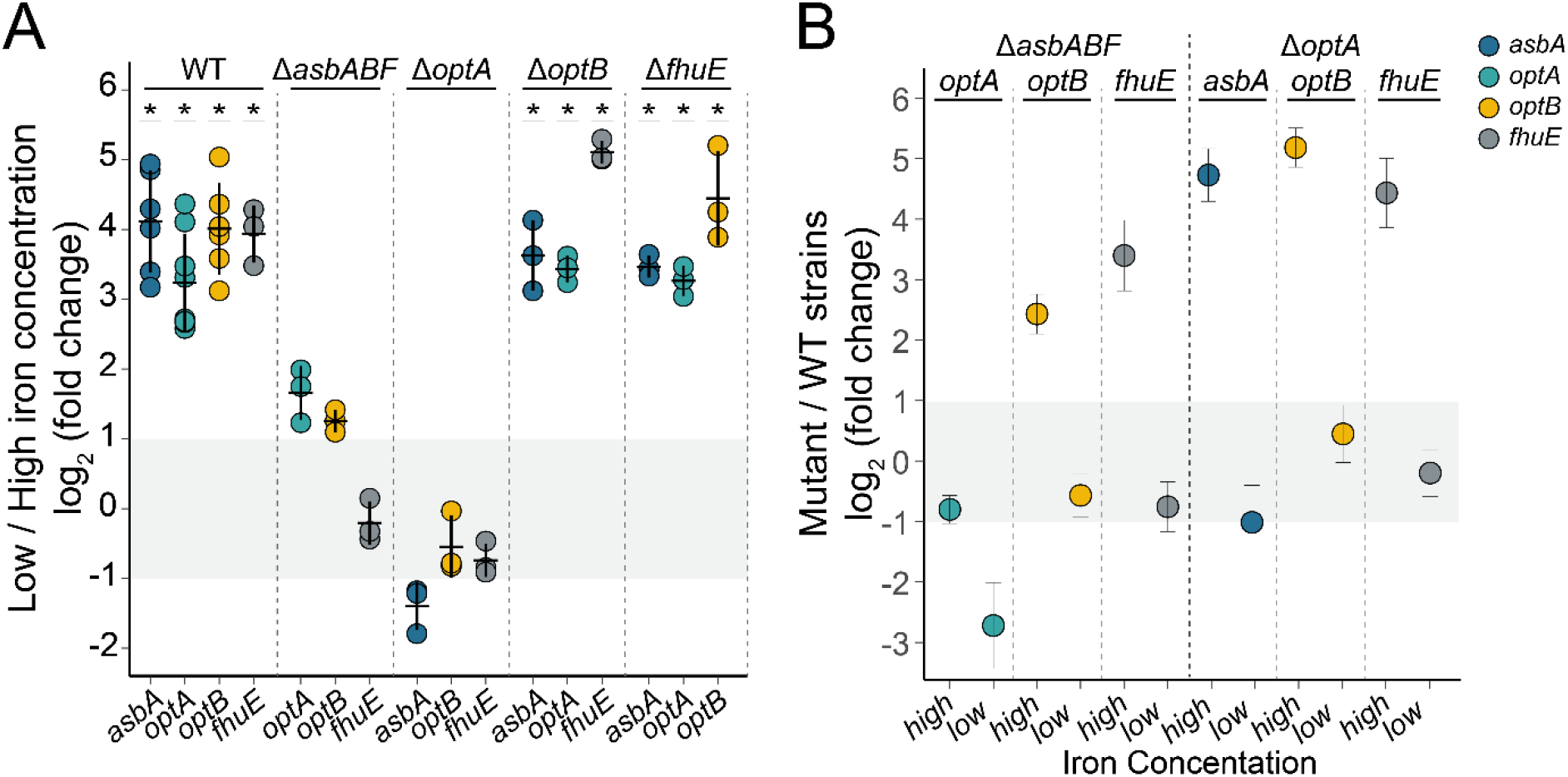
R**e**ciprocal **cross-regulation between *optA* and petrobactin biosynthesis genes.** Results are shown as dot plots based on data from at least three biological replicates (filled circles). RT-qPCR reactions for each sample were carried out in triplicate and normalised to *rpoD* gene expression. Relative fold changes in gene expression were calculated using the 2^-ΔΔCt^ method. (**A**) Expression ratios of *asbA*, *optA*, *optB* and *fhuE* under low versus high iron concentrations in WT and single-mutatnt derivative strains. log_2_ ratios of the expression levels measured by RT-qPCR at 200 nmol·L⁻¹ iron relative to 7 200 nmol·L⁻¹ iron. Error bars represent mean ± SD. Values for the WT from Fig. 2 are shown for comparison. Asterisks indicate log_2_ fold change values significantly different from 1, absence of an asterisk indicates values not significantly different from 1, 0 or –1 (*p* < 0.05, one-sample t-test). (**B**) Expression ratios of *asbA*, *optA*, *optB* and *fhuE* in Δ*asbABF* or Δ*optA* versus WT. log₂ ratios of expression levels measured by RT-qPCR in *ΔasbABF* or Δ*optA* strains relative to the WT. Because mutant and WT biological replicates could not be paired, results are presented as dot plots showing the log₂ ratio of the mean expression of three biological replicates in the mutants to the mean expression of three biological replicates in the WT (individual dots). The SD was estimated from the distribution of fold changes obtained by 2 000 bootstrap resampling iterations, and error bars represent mean ± SD. WT (JM1), Δ*asbABF* (FH35), Δ*optA* (KD17), Δ*optB* (FH42), Δ*fhuE* (FH43).

The loss of iron-responsive regulation observed in the Δ*asbABF* and Δ*optA* strains (**Fig. 4A**) may result from constitutive expression at either high or low levels. To distinguish between these two possibilities, we measured the expression ratio in the Δ*asbABF* and Δ*optA* mutants relative to the wild type, under both low– and high-iron conditions (**Fig. 4B**). The induction of gene expression in the WT can be interpreted as high expression under low-iron conditions and low expression under high-iron conditions (**Fig. 4A**). Therefore, comparing mutant expression to the WT under a given iron condition reveals whether the mutant’s expression level is relatively high or low.

We first examined the effect of the Δ*asbABF* mutation on *optA* expression. Under high-iron conditions, *optA* expression in the Δ*asbABF* was similar to that of the WT strains, remaining at low level (**Fig. 4B**). However, under low-iron conditions, *optA* expression was lower in the Δ*asbABF* mutant than in the WT (**Fig. 4B**). As *optA* expression in the WT is high under low iron-conditions (**Fig. 4A**), *optA* expression in the Δ*asbABF* mutant is interpreted as relatively low. These results suggest that in the absence of petrobactin biosynthesis, *optA* expression remains constitutively low. Similar reasoning applied to *optB* and *fhuE* indicates that these genes are constitutively expressed at high levels in the Δ*asbABF* background. These results suggest that *optA* and *optB* are differentially regulated in response to the absence of petrobactin production.

We next examined strains with individual TBDT mutations. Deletion of *optB* or *fhuE* did not affect the expression of iron uptake genes, with levels remaining comparable to the WT (**Fig. S3B**). In contrast, the Δ*optA* strain grown under high-iron conditions exhibited overexpression of *asbA*, *optB*, and *fhuE* relative to the WT, indicating a high expression level (**Fig. 4B**). Under low iron conditions, their expression levels were similar to the WT, and thus remained high. These findings suggest that in the absence of OptA, the petrobactin biosynthesis pathway and the transporters OptB and FhuE are constitutively overproduced and no longer respond to iron levels in the medium. This indicates that OptA plays a pivotal role in regulating iron acquisition and supports the existence of distinct regulatory functions for OptA and OptB, although they transport the same siderophore.

### Southern Ocean prokaryotes utilise Fe-petrobactin

To investigate the response of microbial consortia originating from the SO to Fe-petrobactin, we carried out incubation experiments with resuscitated communities, one originating from surface waters of the Polar Frontal Zone (PFZ, Station 27) and one from the Antarctic Zone (AAZ, Station 47; **Fig. S1**). Prokaryotic abundances at the initial time point of the batch cultures were 2.45 x 10^4^ cells mL^-1^ in the PFZ experiment and 7.80 x 10^4^ cells mL^-1^ in the AAZ experiment, corresponding to 6% and 19% of the *in situ* cell abundances at the respective sites. After 14 days of incubation, abundances reached 1.14 ± 0.3 x 10^5^ cells mL^-1^ and 1.6 ± 1.37 x 10^6^ cells mL^-1^ in the unamended controls of the PFZ and the AAZ experiments, respectively (**Fig. 5A, C**). In the incubations amended with Fe-petrobactin, cell abundances were significantly higher compared to the controls in both experiments (*p* < 0.0001), demonstrating that Fe-petrobactin was utilised as a source of Fe, thereby enhancing prokaryotic growth.

**Figure 5.**
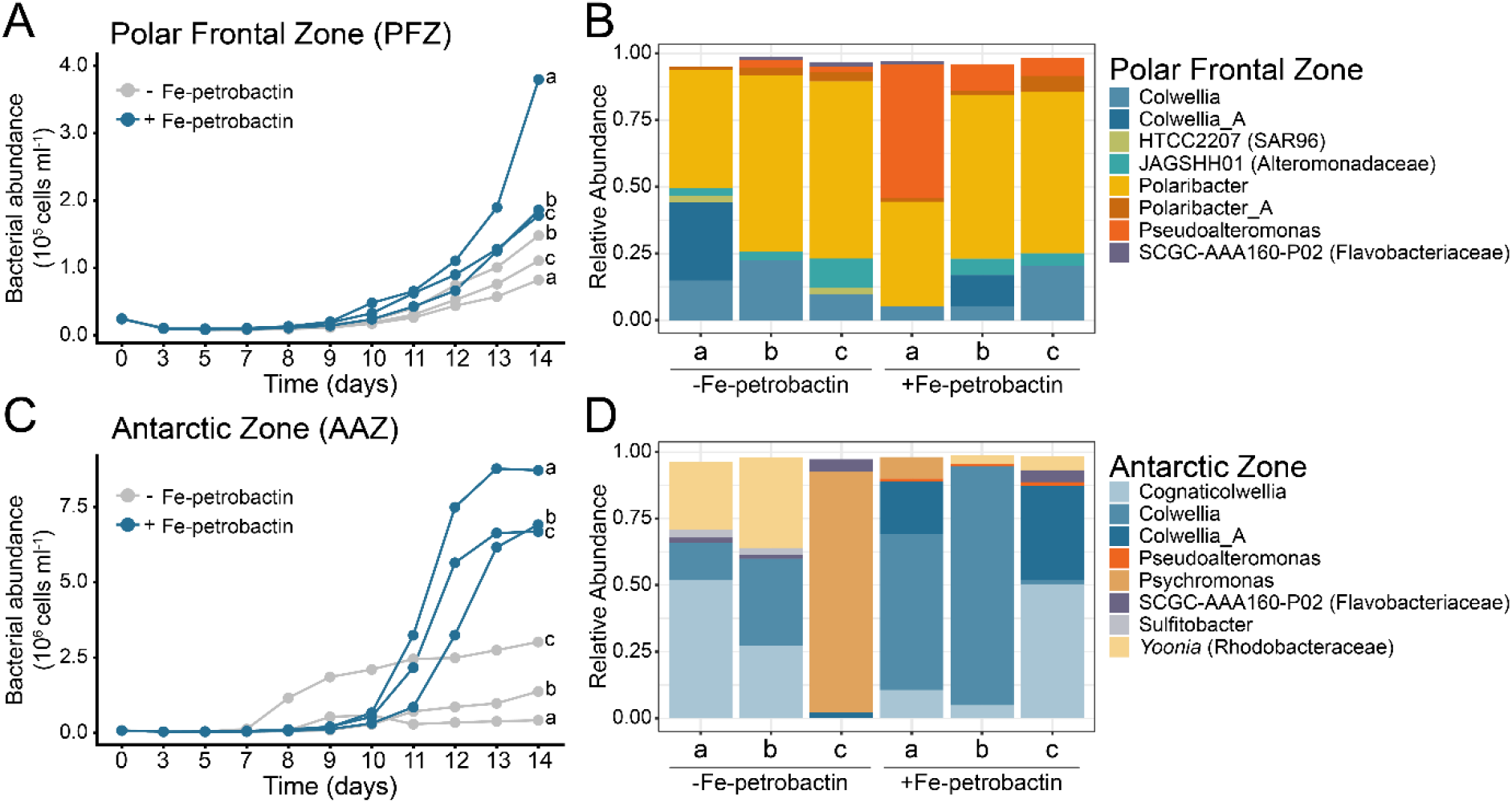
F**e-petrobactin utilisation by resuscitated communities from the Southern Ocean.** Abundances of bacterial cells in batch cultures of resuscitated communities from the Polar Frontal Zone (**A**) and the Antarctic zone (**C**) and composition of the microbial communities at the final time point for the PFZ (**B**) and AAZ (**D**) experiment. Results from triplicate incubations are shown (a-c). The relative abundances of ASVs (>1% in at least one sample) assigned to the genus level in the incubations of the control (ࢤ Fe-petrobactin) and the Fe-petrobactin-amended treatments (+ Fe-petrobactin) are shown. Genus-level assignments followed the GTDB taxonomy and ASVs unresolved to genus level are reported at the deepest confidently classified rank, with the family name given in parentheses. *This incubation most likely represent an outlier regarding the higher growth and the dominance by *Psychromonas* genera (93%).

The composition of the microbial communities differed markedly between the two experiments at the end of the incubation period; however, no significant treatment effect was observed (**Fig. 5B, D**; Bray-Curtis distances and ANOSIM, *p* > 0.5). Only a few taxa exhibited relative abundances that differed between the control and Fe-petrobactin-amended treatments. In the incubations inoculated with the resuscitated communities from the PFZ, one ASV affiliated with *Polaribacter* (Flavobacteriaceae) and two ASVs belonging to *Colwellia* (Alteromonadaceae) had major contributions in all incubations. The cumulative relative abundances of the two *Colwellia* ASVs were overall higher in the control (4-10%) as compared to the Fe-petrobactin-amended incubations (2-3%). The opposite pattern was observed for one ASV belonging to *Pseudoalteromonas* (*Alteromonadaceae*), with a higher share (7-52%) in the incubations amended with Fe-petrobactin as compared to the control (1–3%). The replicate that had the highest contribution of *Pseudoalteromonas* also had the highest cell abundance at the end of the incubation period, suggesting that Fe-petrobactin could have favoured the growth of this taxon. In the AAZ experiment, three ASVs affiliated with *Colwellia* had high relative abundances in all incubations, except for one control. In the Fe-petrobactin-amended treatments, their cumulative abundances reached 60-91%, indicating that these taxa likely benefited from the addition of Fe-petrobactin.

### Fe-petrobactin stimulates taxa harbouring OptB-like transporters

We further screened the metagenomes obtained at the final time point of both incubation experiments for the presence of petrobactin biosynthesis and transport genes. The *asbF* gene, which is specific to petrobactin synthesis, was not detected, indicating the absence of petrobactin biosynthesis in the communities growing in the incubations. Several other siderophore synthesis genes were detected and their GPM varied between experiments, but not between treatments (**Fig. S4A**). The most prevalent siderophore transporter was *pirA* and its relative abundance did not vary between treatments or experiments (**Fig. S4B**). To explore siderophore synthesis and transport at the taxon level, we profiled 62 high-quality MAGs (>80% completion, <10% contamination; **Table S9**). Gene clusters encoding the synthesis of the siderophores pyochelin (*pch*), rhizobactin (*rhb*) and vanchrobactin (*vab*) were detected in three Alteromonadaceae-affiliated MAGs (MAG15, MAG64, MAG81). Siderophore transport genes were found in 36 MAGs with the *pir*A gene being the most widespread (**Fig. S5**).

Given the prevalence of the *pirA* gene, which is known to encode a general transporter, we investigated whether this gene could potentially facilitate the transport of petrobactin. We performed MUSCLE global alignments comparing PirA sequences detected in the metagenomes and MAGs from our experiments with the petrobactin transporter sequences OptA and OptB from *M. nauticus* (**Table S10**). A total of 106 and 246 *pirA* genes were detected in the metagenomes from the PFZ and the AAZ experiments, respectively. We found that none of the encoded protein sequences showed more than 20% amino acid (AA) identity with OptA. (data not shown), but we identified four with high AA identity to OptB. The two genes encoding proteins with the highest AA identity to OptB (49.9% and 51.2%) were taxonomically affiliated with *Pseudoalteromonas* and exhibited significantly higher relative abundances (GPM) in the Fe-petrobactin treatments compared to the control in both experiments (**Fig. 6A**). Furthermore, MUSCLE global alignments across the 62 MAGs confirmed that no PirA sequences showed high similarities to the OptA from *M. nauticus* (< 22.5%) (**Table S10**). However, two MAGs affiliated with Alteromonadaceae, specifically *Pseudoalteromonas*, encode PirA sequences with 54.6% (MAG15) and 53.7% (MAG64) AA identity to OptB.

**Figure 6.**
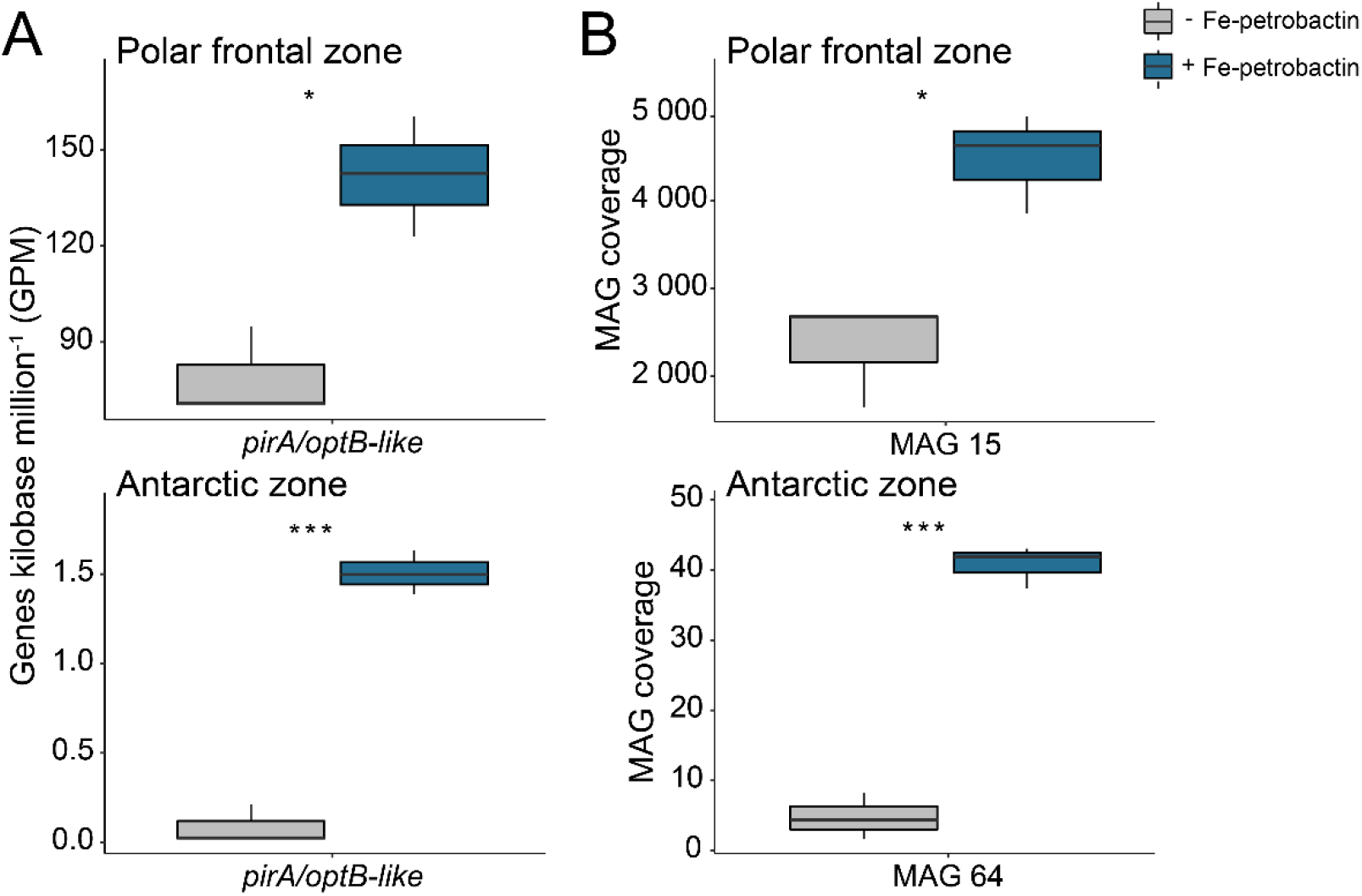
M**A**G15 **and MAG64 respond specifically to Fe-petrobactin amendment.** (**A**) Relative abundance (genes per kilobase million, GPM) of *pirA*/*optB*-like genes (> 49.9% amino acid similarity with the OptB transporter in *M. nauticus*) in the metagenomes of the Polar Frontal Zone and Antarctic Zone experiments. (**B**) Coverage of MAGs containing the *pirA*/*optB*-like genes in the Polar Frontal Zone and Antarctic Zone experiments. MAG15 and MAG64 are taxonomically affiliated with *Pseudoalteromonas*. Mean of triplicate incubations ± SD are shown. Grey: control experiment; dark bue: petrobactin amended treatment. Significant differences among treatments are indicated (\**p* < 0.05, \*\**p* < 0.01, \*\*\**p* < 0.001).

Hereafter, PirA proteins sharing high AA identity to OptB from *M. nauticus* (30-50%) are referred to as OptB-like. The coverage of these two MAGs in the metagenomes was significantly higher in the Fe-petrobactin–amended incubations compared to the controls (**Fig. 6B**), a pattern that was not observed for any other MAG (**Fig. S6**). Taken together, our results on the metagenome and MAG levels demonstrate that taxa encoding OptB-like transporters respond specifically to Fe-petrobactin addition in our incubations.

We further explored whether *optB*-like transporter genes were present in marine microbial communities, by examining 23 (< 0.8 µm size fraction) and 19 (> 0.8 µm size fraction) metagenomes, as well as 556 MAGs reconstructed from samples collected throughout the water column along the GEOTRACES SWINGS transect ([47]; **Fig. S1**). We detected 26 OptB-like sequences (30% to 56% AA identity) encoded in the metagenomes and in 8 MAGs (32% to 59% AA identity; **Table S10**). The OptB-like transporters in the metagenomes were taxonomically affiliated with Pseudomonadaceae, Halomonadaceae, Marinomonadaceae, SAR86 and *Alteromonas*. The MAGs were assigned to the Alteromonadaceae (*Pseudoalteromonas*, *Alteromonas*), Halomonadaceae (*Halomonas*, *Halopseudomonas*), and Sphingomonadaceae. The samples from which the metagenomes and MAGs were reconstructed originated mainly from deep water masses across the transect (**Fig. S7**).

### OptA and OptB have different evolutionary history

To investigate the evolutionary relationships of the petrobactin transporters identified in our study, we performed a phylogenetic analysis using putative petrobactin transporters retrieved from the Integrated Microbial Genomes (IMG) database. Putative petrobactin transporters were identified as TBDTs located in the genomic vicinity of predicted petrobactin-related biosynthetic gene clusters. These clusters contain genes that are annotated as spermidine-citrate ligase, petrobactin synthase and 3-dehydroshikimate dehydratase (see Supplementary Information). Putative petrobactin biosynthetic clusters were identified in *Rhodopseudomonas palustris* CGA009 and *Phaeobacter inhibens* 2.10 which produce rhodopetrobactin A and B and roseobactin, respectively. These siderophores share structural similarities with petrobactin, as they are assembled from spermidinyl (or homospermidinyl), citryl, and 3,4-dihydroxybenzoyl moieties (**Fig. 7**). This structural similarity supports the accuracy of the search for biosynthesis gene clusters responsible for producing petrobactin-related siderophores. The analysis also included a set of TBDTs for which the transported siderophores had been determined experimentally (**Table S11**).

**Figure 7.**
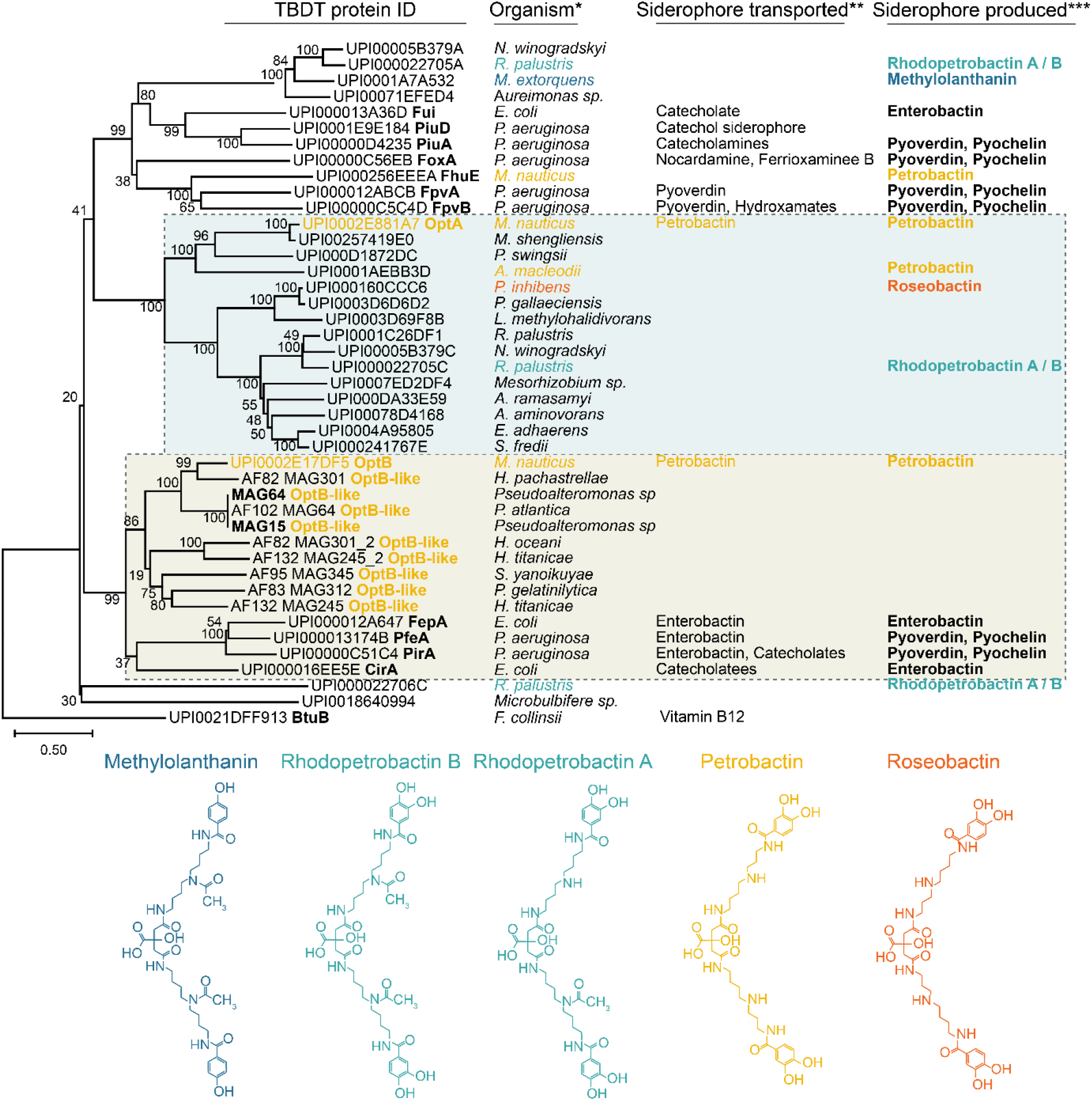
P**h**ylogenetic **tree of petrobactin related TBDTs.** The phylogenetic tree, generated using the neighboour-joining method is drawn to scale, with branch lengths representing the number of substitutions per site. The Vitamin B transporting TBDT BtuB from *F. collinsii* was added as a subgroup. Node support was assessed by bootstrap analysis with 2,000 replicates. UniParc ID was used for sequences collected from databases. Protein names were added when known. OptA, OptB and OptB-like proteins are indicated in yellow. Coloured backgrounds indicate the two clusters that include OptA (blue) and OptB (yellow) sequences from *M. nauticus*. Structures of petrobactin (yellow) and petrobactin-related siderophores (methylothanin in blue, rhodopetrobactin in green and roseobactin in orange) are shown below the tree. *Full name of the strains and references can be found in **Table S11**; **Siderophores are indicated when there is evidence of transport by the corresponding TBDT in the literature; ***Siderophores are indicated when there is evidence of production by the corresponding strain in the literature.

OptA groups with TBDTs associated with predicted petrobactin biosynthesis genes, whereas OptB clusters with homologs from MAGs derived from SO communities, forming two well-supported and distinct clades. This phylogenetic separation suggests that OptA-like and OptB-like TBDTs followed distinct evolutionary trajectories. The OptA family likely co-evolved with petrobactin production pathways, whereas the OptB family seems to have evolved independently.

## Discussion

### OptA and OptB mediate Fe-petrobactin uptake

Although TBDTs are readily detected in marine genomes and metagenomes, identifying their substrates based on sequence analysis remains challenging and continue to be a bottleneck in the mechanistic understanding of siderophore production and uptake in marine environments. In this work, we demonstrate that OptA and OptB are two transporters of Fe-petrobactin in *M. nauticus*. Despite transporting the same siderophore, OptA and OptB share only 24% sequence identity, indicating that low sequence identity does not preclude the sharing of a common substrate. A BLAST search for OptA homologs in *Alteromonas macleodii*, which produces petrobactin, gave MASE_17930 as the best hit (37% identity), suggesting it could mediate petrobactin uptake in this bacterium. This is further supported by previous gene neighbourhood analysis and phylogenetic distribution across the *Alteromonas* genus that led Manck et al. [48] to propose MASE_17930 as the petrobactin transporter. Nevertheless, unlike OptA, MASE_17930 is not co-located with the petrobactin biosynthesis gene cluster. These observations demonstrate that sequence similarity and gene neighbourhood alone cannot reliably predict the substrate specificity of TBDTs, underscoring the need for experimental validation when annotating TBDTs in marine bacteria. Moreover*, A. macleodii* appears to lack a TBDT closely related to OptB, suggesting that strategies for petrobactin uptake may vary among bacterial taxa.

The genomic organization and functional redundancy of *optA* and *optB* in *M. nauticus* are comparable to those of *fpvA* and *fpvB*, which encode transporters of Fe-pyoverdine in *Pseudomonas aeruginosa*. Specifically, *optA* and *fpvA* are adjacent to their respective siderophore biosynthetic gene clusters, whereas *optB* and *fpvB* are positioned elsewhere on the chromosome [49]. Although both FpvA and FpvB function as receptors for Fe-pyoverdine, they have different substrate specificity. FpvA shows strict specificity for Fe-pyoverdine, whereas FpvB displays broader ligand recognition, accommodating structurally diverse substrates through distinct binding modes [50,51]. By analogy, OptA may functions as a primary, dedicated transporter, while OptB could act as a more promiscuous secondary receptor.

González et al. [52] demonstrated that the secondary Fe-pyoverdine transporter, FpvB, is advantageous under certain environmental conditions but detrimental under others. It is therefore conceivable that the presence of OptA and OptB provides a versatile strategy for Fe-petrobactin uptake, optimizing the cost-benefit ratio across the diverse environmental conditions encountered in marine habitats.

### A versatile petrobactin uptake strategy

RT-qPCR experiments demonstrated that the TBDT genes *optA*, *optB*, and *fhuE*, as well as the petrobactin biosynthesis gene *asbA*, are upregulated under low-iron conditions. Transcriptional regulation of iron-uptake genes in response to iron availability is typically mediated by the ferric uptake regulator Fur [53]. It is therefore likely that a similar regulatory mechanism operates in *M. nauticus*. However, the iron-responsive regulation is lost in the Δ*asbABF* and Δ*optA* mutants. This suggests that a regulatory mechanism dependent on the presence of OptA and petrobactin contributes to the modulation of iron-uptake gene expression. In contrast, OptB, does not appear to participate in this regulation, as the iron-responsive regulation remains effective in a Δ*optB* mutant (**Figure 4A**, **Figure S3B**). These findings indicate that although OptA and OptB transport the same siderophore they are not functionally equivalent regarding their regulatory roles.

Comparisons of expression levels in Δ*asbABF* and WT under high– and low-iron conditions revealed that, in the absence of petrobactin biosynthesis, *optA* expression is constitutively low, while *optB* and *fhuE* expression are constitutively high. This suggests that in the absence of petrobactin, iron transport by OptB and FhuE may be favoured. Since FhuE does not transport petrobactin, its increased expression may allow the uptake of alternative Fe-siderophores, thereby compensating for the defect in petrobactin-mediated iron acquisition. The similar regulatory pattern observed for the *optB* gene, with increased expression in the absence of petrobactin production, supports the hypothesis that OptB could also transport other siderophores.

OptA clusters within a clade that comprising TBDTs whose genes are located near predicted petrobactin biosynthesis gene clusters, suggesting that these transporters may have co-evolved with petrobactin biosynthesis pathways and likely function as their primary transporters. In contrast, OptB and OptB-like TBDTs from MAGs are not located near a petrobactin biosynthesis gene cluster and form a distinct group. This phylogenetic separation of OptB transporters from OptA hints at a different evolutionary history, potentially independent of petrobactin synthesis. OptB may have followed an alternative trajectory, independent of the ability to synthesize petrobactin, and may have instead evolved the ability to import petrobactin as a xenosiderophore.

### OptB-like transporters could favour the utilisation of Fe-petrobactin in marine microbial communities

Fe-petrobactin led to enhanced growth of Southern Ocean bacteria in carbon-replete incubations, favouring taxa harbouring OptB-like transporters (**Fig. 6**). Our incubations were carried out in seawater collected from SO surface waters, which likely contained Fe bound to various siderophores, some of which might be recognized by OptB-like transporters. However, given that the concentration of Fe-petrobactin in our incubations exceeded that of individual siderophores present in surface seawater [7–9], the observed response was likely driven by the addition of this specific Fe form. In this study, the *optB*-like genes were initially identified as belonging to the *pirA* group, which is widespread in marine microbial communities [9,47] and considered as encoding a general transporter. Our results suggest that a subset of PirA proteins may function as OptB-like transporters, potentially facilitating petrobactin-mediated Fe uptake.

The marked differences in community composition between the two experiments could be explained by distinct resident microbial communities in the respective sampling regions [54] and by the taxa capable of grow upon resuscitation. Based on the relative abundances of the ASVs and the taxonomic affiliation of *optB*-like genes at the metagenome and MAG level, *Pseudoalteromonas* and *Colwellia* were the most responsive to Fe-petrobactin additions. Both are ubiquitous marine bacteria genus known to possess mechanisms of adaptation to cold temperatures [55,56]. Their diverse repertoire of siderophore-associated functional capabilities [13,57,58] could have been advantageous under our experimental conditions, where abundant labile organic carbon supported their copiotrophic lifestyle [59]. Two environmental MAGs harbouring OptB-like transporters were affiliated with *Pseudoalteromonas* (**Table S10**), suggesting that this genus could play a role in Fe-petrobactin acquisition under varying conditions. OptB-like transporters identified *in situ* were further affiliated with other taxonomic groups such as Pseudomonadaceae, Halomonadaceae and *Alteromonas* known for their siderophore-producing and transport capacities [3,8,60]. The absence of the *asbF* gene, essential for petrobactin synthesis, and of potential *optA*-like genes indicate that Fe-petrobactin is used as a xenosiderophore by these taxa. The recurrence of OptB-like transporters across the SO extends our experimental observations to a broader context, suggesting that this uptake mechanism could be a specialised strategy for some microbial community members. Transporters that exploit exogenous siderophores enable cells to benefit from siderophores produced by others while avoiding the energetic costs of biosynthesis. Such dynamics reflect well-documented “cheating” or siderophore piracy behaviours [16,17,61,62] with important implications for Fe cycling.

Taken together, the integration of model organisms, genetic tools, community-level responses, genome-resolved analyses on microbial communities from incubation experiments and *in situ* observations highlights a previously underappreciated strategy, suggesting that OptB-like transporters could contribute to Fe uptake in the ocean.

## Acknowledgements

We would like to thank Corentin Baudet and Antoine Ringard for providing surface seawater using the GeoFISH, Kilian Ducos for the first qPCR experiments and François Rigal for advices on the phylogenetic tree and some statistical tests. To detect typos and improve the flow of the English in the manuscript, an artificial intelligence tool based on a large language model (LLM) was used.

## Author contributions

**SN**: Conceptualization, Investigation, Methodology, Supervision, Validation, Visualization and Writing—review & editing. **RT**: Data curation, Investigation, Methodology, Validation, Visualization and Writing – original draft. **FH**: Investigation, Methodology and Validation. **AG**: Investigation, Methodology and Validation. **PC**: Investigation, Methodology and Validation. **KK**: Investigation, Methodology and Validation. **RL**: Funding acquisition, Reviewing. **SB**: Conceptualization, Funding acquisition, Investigation and Writing—review & editing. **IO**: Conceptualization, Funding acquisition, Validation, Investigation and Writing— review & editing. **RG**: Conceptualization, Funding acquisition, Investigation, Methodology, Supervision, Validation, Writing—original draft and Writing—review & editing

## Data availability

Raw sequence data were deposited to the European Nucleotide Archive (ENA) under the project accession PRJEB873833 for 16S rDNA amplicons and the project accession PRJEB86053 for the metagenomics samples.

## Supplementary data

**Supplementary Information** (pdf file) File includes the supplementary material and methods, Tables S1 to S4; S6 to S9 and S11, Figures S1 to S7 and References.

**Table S5** (.xlsx file) List of all amplicon sequence variants (ASVs) with taxonomic assignments and respective relative abundances ordered based on the type of treatments (control and Fe-petrobactin amended) for the Antarctic Zone and Polar Frontal Zone experiments.

**Table S10** (.xlsx file) List of protein sequences from MAGs and metagenomes used for MUSCLE global alignment against reference sequences OptA or OptB.

