## SupplementaryInformations for "TonB-dependent Transporters of Fe-petrobactin in *Marinobacter nauticus* and Southern Ocean bacteria"

##### File content:

- Supplementary Materials and Methods
- Tables S1 to S4, S6 to S9 and S11
- Figures S1 to S7
- References

#### Supplementary Materials and Methods

##### Purification of petrobactin and preparation of Fe-petrobactin complexes

Petrobactin was purified as describe in Kińska et al. [1]. Fe-petrobactin complexes were prepared from purified petrobactin. For complex formation, 60 µL of petrobactin solution at 650 µM in water, 80 µL of ammonium acetate 100 mM, pH 8-9, 20 µL of 100 mg $L^{-1}$ Fe-citrate and 240 µL type 1 deionized water (MilliQ) were mixed. To ensure quantitative iron binding to petrobactin, complexes were allowed to form overnight. Complex formation was confirmed by separation on an Acquity UPLC Protein BEH SEC column (4.6 x 150 mm, Waters) connected to an Agilent 1200 LC system with 10 mM ammonium acetate, pH 7.9-8.0, as a mobile phase (15 min, 0.3 mL $min^{-1}$, 25°C) with elemental detection by ICP-MS with an Agilent 7700X ICP-MS analyzer, using hydrogen at 4 mL min-1 as reaction gas to suppress spectral interferences. Fe-petrobactin complexes were purified from Fe-citrate on a Superdex® Peptide 10/300 GL SEC column (separation range 100 and 7000 Da, 10 × 300 mm, GE Healthcare) using 10 mM ammonium acetate pH~8 as mobile phase. Fractionation was monitored based on the UV-Vis signal at 480 nm, which coincided with the elemental signal, as verified for previous samples. For each sample 3 fractions were collected and subsequently their purity was checked on UPLC column with elemental detection. The fractions with the highest purity were used in further experiments.

**RT-qPCR after supplementation of cultures with petrobactin**

Growth conditions used are similar to the supplementation of TBDT mutant with petrobactin experiment. *M. nauticus* strains were streaked from -80°C glycerol stock onto fresh EMS (7.2 µM $FeCl_2$) plates containing 20 mM lactate and 100 µM streptomycin and incubated at 30°C for five days. A few colonies were then resuspended in 0.5 mL of iron-free MEM medium. The cell suspension was diluted appropriately in iron-free MEM medium, before the inoculation, in triplicates, at an $A_{600nm}$ of 0.1 in MEM medium supplemented with 1 µM $FeCl_3$, 200 µM EDTA and 10 µM of petrobactin when indicated. After 16 h of incubation, 2,4 ml of cells were harvested and resuspended in 200 µL of 10 mmol $L^{-1}$ Tris-HCl pH8. RNAs were extracted as previously described [2], using the Extract-All (Eurobio Scientific) kit, then subjected to DNase treatment using the TURBO DNA-free™ kit (Invitrogen). 325 ng of RNA were used to prepare cDNAs using the SuperScript™ III First-Strand Synthesis SuperMix for qRT-PCR kit (Invitrogen). Expression of specific genes was measured in a CFX connect system (Biorad) with Bio-Rad CFX Maestro software using the Luna® Universal qPCR Master Mix (NEB) with the appropriate primers (**Table S3**). qPCR reactions were prepared in duplicates, in a 96 wells PCR plates, using the epMotion® 5070 (Eppendorf). 1,25 µg of cDNA were used for each reaction.

**Bioinformatics pipeline**

The short reads from the six samples for each experiment that passed the quality control (Table S6) were co-assembled using *megahit* v1.2.9 [3] with a minimum length of 1000 bp. For comparison, *metaspades* v3.15.5 was also used for the assembly of the short reads. Based on a comparison using *quast* v.5.0.2 [4], *megahit* was used for further analysis. To develop a non-redundant database, the contig file was reformatted with *anvi'o* [5] using the parameter --min-len 2000. *Prodigal* v2.6.3 was used to predict open reading frames (ORFs) [6] with the parameters *-p meta*. *Salmon* v.1.10.2 [7]was later used to quantify each gene in each sample and to assign short reads to the non-redundant gene sequences in the database with parameters set at *--meta*. The quantified occurrence of each gene in each sample was normalized as a metric of genes per kilobase million (GPM) for the metagenome to eliminate the library size effect and different gene lengths [8,9]. This ensured the comparability between the samples. To retrieve the GPM for each identified protein at 100 % similarity, the protein sequence corresponding to a gene related to the respective siderophore pathway in each sample was merged with gene coverage data obtained using *anvi'o* [5] with the parameter *anvi-profile-blitz* combined with salmon output data.

The tool *FeGenie* [10] was used to detect genes with Fe-related pathways based on separate contig files for the subsequent alignment and annotation of each gene detected. This tool uses Hidden Markov Models (HMMs) along with blast to annotate iron-related genes. In this study, we used *FeGenie* to classify the proteins related to siderophore synthesis and transport. In the existing dataset, the identified genes passed the predetermined bit score cut-offs. Although not all subfamilies of each gene family in each sample were identified, it must be noted that the HMMs developed in *FeGenie* were sensitive to the presence of the entire gene family. One of the reasons for this could be the low GPM of genes related to siderophore synthesis and transport. The GPM data were combined with the FeGenie results to compute statistical comparisons between the -Fe-petrobactin and +Fe-petrobactin amended samples using two-sample t-tests, and a statistically significant result was considered at $p < 0.05$. Significant siderophore synthesis and transport gene families were taxonomically screened using BLAST+ [11].

**Genome-resolved metagenomics**

For effective binning, the *anvi'o* pipeline was used [5]. Short reads from the co-assembled metagenomes were mapped to the scaffolds using *bowtie2* v2.5.1 [12] and stored as BAM files with *samtools* v.1.15.1 [13] with a minimum identity of 95 %. Using the *anvi-init-bam* option from *anvi'o*, the alignment files of each sample were indexed and used to reconstruct metagenome-assembled genomes (MAGs). Because the metagenomes were co-assembled, we expected the highest completion of MAGs. Using *anvi'o*, each binary alignment/map (BAM) file was profiled to obtain the coverage and detection statistics for each scaffold to create combined mapping profiles, which were then merged into a profile database. Contigs were manually binned using *MetaBAT* v.2.15 [14], *MaxBin* v.2.2.7 [15], and *Concoct* v.1.10 [16]to ensure comparability between each binning tool. To retrieve a non-redundant set of bins, *DAS_Tool* v1.1.3 [17]was used to dereplicate, aggregate, and score each MAG. The resulting MAGs were imported into *anvi'o* for refinement based on their GC content, differential coverage, and sequence composition. Later, taxonomic identification of MAGs was performed internally using *anvi'o*. The relative abundance of each MAG was determined by assessing the percentage of reads recruited from each MAG across all the samples. *CheckM* v1.2.2 [18]and *anvi'o* were used to reassess each MAG for high completion and low redundancy statistics (**Table S9**). To elucidate siderophore-related pathways, the *antiSMASH* tool [19] was employed to identify secondary metabolite biosynthetic gene clusters (BGCs) associated with siderophore biosynthesis and transport. Notably, the *antiSMASH* tool does not show specific gene transporters for siderophores but rather common transporters for secondary metabolite biosynthesis gene clusters, which may include transport-related genes for siderophores [20]. These results were combined with the results of the FeGenie that was also conducted on the final MAGs to obtain siderophore-related gene information on each MAG along with known taxonomy. Additionally, a multiple sequence global alignment tool like *MUSCLE* (Multiple Sequence Comparison by Log-Expectation) was implemented to compare siderophore-related genes in our MAGs of interest compared to the reference gene sequences of the strains using a pairwise distance estimation based on k-mer similarity [21].

**TBDT genes retrieval for phylogenetic analysis**

Genomes containing *asbF* homologs located within a NIS-type siderophore biosynthetic gene cluster were identified by Manck et al. [22] as potential petrobactin biosynthesis clusters. These genomes were retrieved from the Integrated Microbial Genomes (IMG) database [23]. To refine the dataset, genomes classified within the Bacillota phylum were removed, and only genomes with the highest sequence quality (Level 6: Finished) were retained, resulting in a final set of 93 genomes. The cassette search tool provided by the IMG/M system was used to identify gene clusters containing genes from the following families: pfam04183 (IucA/IucC family), pfam19468 (biosynthetic gene cluster family), and genes encoding a putative 3-dehydroshikimate dehydratase (EC 4.2.1.118), with a maximum allowed distance of 5 kb between query genes. To capture the surrounding genomic context, 10 kb were added to each end of the retrieved regions. From the cassette search, 71 gene clusters were retrieved. Genes encoding putative TBDTs (pfam00593) were then searched within the cassettes. Of these, 53 cassettes contained one putative TBDT gene, 13 cassettes did not contain any, and five cassettes contained 2 or 3 TBDT genes each. In total, 65 putative TBDT genes were retrieved.

**Table S1 Bacterial strains used in this study.**

[a] The following abbreviations refer to resistance to: kanamycin ($Kn^R$), Nalidixic acid ($Nal^R$), trimethoprim ($Tp^r$) and Streptomycin ($Sm^R$).

| Strain | Relevant genotype[a] | Source or reference[a] |
|---|---|---|
| ***Escherichia coli*** | | |
| **S17-1 (λpir)** | F-, RP4-2(*Km*::Tn*7*,Tc::Mu-1), *pro-82*, *λpir*, *recA1*, *endA1*, *thiE1*, *hsdR17*, *creC510* ($Tp^R$, $Sm^R$) | Donor strain for conjugation. [24] |
| **DH5α (λpir)** | supE44, ΔlacU169 (ΦlacZΔM15), recA1, endA1, hsdR17, thi-1, gyrA96, relA1, λpir | [25] Host for cloning. |
| **PIR1** | F- Δ*lac169 rpoS(Am) robA1 creC510 hsdR514 endA recA1 uidA*(Δ*MluI*)::*pir-116* | Invitrogen |
| **FH32** | DH5α (λpir) / pFH32 | pFH32 × DH5α (λpir) to $Kn^r$ |
| **FH33** | DH5α (λpir) / pFH33 | pFH33 × DH5α (λpir) to $Kn^r$ |
| **FH37** | S17-1 (λpir) / pFH32 | Transformation pFH32 × S17-1 (λpir) to $Kn^r$ |
| **FH38** | S17-1 (λpir) / pFH33 | Transformation pFH33 × S17-1 (λpir) to $Kn^r$ |
| **FH39** | S17-1 (λpir) / pSN81 | Transformation pSN81 × S17-1 (λpir) to $Kn^r$ |
| **FH74** | PIR1 / pFH74 | pFH74 × PIR1 to $Kn^R$ |
| **FH75** | PIR1 / pFH75 | pFH75 × PIR1 to $Kn^R$ |
| **FH76** | PIR1 / pFH76 | pFH76 × PIR1 to $Kn^R$ |
| **FH77** | S17-1 (λpir) /pFH74 | Transformation pFH74 × S17-1 (λpir) to $Kn^R$ |
| **FH78** | S17-1 (λpir) /pFH75 | Transformation pFH75 × S17-1 (λpir) to $Kn^R$ |
| **FH79** | S17-1 (λpir) /pFH76 | Transformation pFH76 × S17-1 (λpir) to $Kn^R$ |
| **KD36** | S17-1 (λpir) /pSN81 | Transformation pSN81 × S17-1 (λpir) to $Kn^R$ |
| ***Marinobacter nauticus*** | | |
| **SP17** | ATCC 49840 | [26] |
| **JM1** | SP17 *rpsl*K58T ($Sm^R$) | [27] |
| **FH35** | JM1 Δ*asbABF* | [2] |
| **KD17** | JM1 Δ*optA* | Conjugation KD36 × JM1 to $Kn^R$ $Sm^S$ and allelic exchange to $Kn^S$ and $Sm^R$ |
| **FH42** | JM1 Δ*optB* | Conjugation FH37 × JM1 to $Kn^R$ $Sm^S$ and allelic exchange to $Kn^S$ and $Sm^R$ |
| **FH43** | JM1 Δ*fhuE* | Conjugation FH38 × JM1 to $Kn^R$ $Sm^S$ and allelic exchange to $Kn^S$ and $Sm^R$ |
| **FH46** | JM1 Δ*asbABF* Δ*optA* | Conjugation FH39 × FH35 to $Kn^R$ $Sm^S$ and allelic exchange to $Kn^S$ and $Sm^R$ |

| | | |
|---|---|---|
| **FH50** | JM1 Δ*asbABF* Δ*optB* | Conjugation FH37 × FH35 to $Kn^R$ $Sm^S$ and allelic exchange to $Kn^S$ and $Sm^R$ |
| **FH51** | JM1 Δ*asbABF* Δ*fhuE* | Conjugation FH38 × FH35 to $Kn^R$ $Sm^S$ and allelic exchange to $Kn^S$ and $Sm^R$ |
| **FH71** | JM1 Δ*asbABF* Δ*optA* Δ*optB* | Conjugation FH37 × FH46 to $Kn^R$ $Sm^S$ and allelic exchange to $Kn^S$ and $Sm^R$ |
| **FH72** | JM1 Δ*asbABF* Δ*optA* Δ*fhuE* | Conjugation FH38 × FH46 to $Kn^R$ $Sm^S$ and allelic exchange to $Kn^S$ and $Sm^R$ |
| **FH80** | JM1 Δ*RS09625* | Conjugation FH79 × JM1 to $Kn^R$ $Sm^S$ and allelic exchange to $Kn^S$ and $Sm^R$ |
| **FH81** | JM1 Δ*btuB* | Conjugation FH77 × JM1 to $Kn^R$ $Sm^S$ and allelic exchange to $Kn^S$ and $Sm^R$ |
| **FH82** | JM1 Δ*RS01205* | Conjugation FH78 × JM1 to $Kn^R$ $Sm^S$ and allelic exchange to $Kn^S$ and $Sm^R$ |
| **FH83** | JM1 Δ*asbABF* Δ*optA* Δ*RS09625* | Conjugation FH79 × FH46 to $Kn^R$ $Sm^S$ and allelic exchange to $Kn^S$ and $Sm^R$ |
| **FH84** | JM1 Δ*asbABF* Δ*optA* Δ*btuB* | Conjugation FH77 × FH46 to $Kn^R$ $Sm^S$ and allelic exchange to $Kn^S$ and $Sm^R$ |
| **FH85** | JM1 Δ*asbABF* Δ*optA* Δ*RS01205* | Conjugation FH78 × FH46 to $Kn^R$ $Sm^S$ and allelic exchange to $Kn^S$ and $Sm^R$ |

**Table S2 Plasmids used in this study.**

| Name | Construct and Usage | Source or reference |
|---|---|---|
| **pSN77** | pSEVA211 derivative, carries *rpsL* gene from *M. nauticus* SP17, conferring Streptomycin sensitivity | [2] |
| **pSN81** | pSN77 derivative, carries Δ*optA locus*, used for allelic exchange | Insertion of *optA*-flanking sequences using Restriction/Ligation |
| **pFH32** | pSN77 derivative, carries Δ*optB locus*, used for allelic exchange | Insertion of *optB*-flanking sequences using Prolonged Overlap Extension-PCR (POE-PCR [28]) |
| **pFH33** | pSN77 derivative, carries Δ*fhuE locus*, used for allelic exchange | Insertion of *fhuE*-flanking sequences using POE-PCR |
| **pFH74** | pSN77 derivative, carries Δ*btuB locus*, used for allelic exchange | Insertion of *btuB*-flanking sequences using POE-PCR |
| **pFH75** | pSN77 derivative, carries Δ*RS01205 locus*, used for allelic exchange | Insertion of *RS01205*-flanking sequences using POE-PCR |
| **pFH76** | pSN77 derivative, carries Δ*RS09625 locus*, used for allelic exchange | Insertion of *RS09625*-flanking sequences using POE-PCR |

**Table S3 Primers used in this study.**

| Name | Sequence | Construct |
|---|---|---|
| **OL52** | CAGTCCGCGGCTCCTGGGTGTATTGAC | Amplification of *optA*-flanking sequences for pSN81 construct by digestion and ligation using SacII, EcoRI and SacI |
| **OL53** | GCCAGAATTCATGTTGCGTTACCAAATCAC | |
| **OL54** | ATCTGAATTCGACCGCACTCCATTATGAAA | |
| **OL55** | ATTAGAGCTCCTGGTGGATACCATTGCG | |
| **OL109** | CTTGGACTCCTGTTGATAGATCCAGT | Amplification of pSN77 vector for pFH32, pFH33, pFH74, pFH75 and pFH76 construct by POE-PCR |
| **OL110** | CCATGTCAGCCGTTAAGTGTTCC | |
| **OL139** | AGCTGTTGCTGGTAGAAC | Checking Δ*optA* by the amplification of flanking sequences in FH46, FH71, FH72, FH83, FH84 and FH85 strains |
| **OL140** | CCATCAGTTGCGTAGAGG | |
| **OL181** | ACACTTAACGGCTGACATGGAAATCCGGATGCAGCAGTAC | Amplification of *fhuE*-flanking sequences for pFH33 construct by POE-PCR |
| **OL182** | GACTGCCCCACCCTGTGACTAACGGCTCCTTGGTTG | |
| **OL183** | CAACCAAGGAGCCGTTAGTCACAGGGTGGGGCAGTC | |
| **OL184** | GATCTATCAACAGGAGTCCAAGCAGATGCAAGACTGGTAC | |
| **OL187** | GCCAGAGTCCACTTCCAG | Checking Δ*fhuE* by the amplification of flanking sequences in FH43, FH51 and FH72 strains |
| **OL188** | CAGTTCAAATGCGAGGCC | |
| **OL189** | ACACTTAACGGCTGACATGGCATCATCCCTGGGTGGATCTG | Amplification of *optB*-flanking sequences for pFH32 construct by POE-PCR |
| **OL190** | CATTGAAGTCATGGTTACTATCCGATGTATCAGTTTTCGCATTTGC | |
| **OL191** | GCAAATGCGAAAACTGATACATCGGATAGTAACCATGACTTCAATG | |
| **OL192** | GATCTATCAACAGGAGTCCAAGCTGCCGAAGTGAAAAGTTC | |
| **OL195** | CGGTCATCAGTACCAATCCTG | Checking Δ*optB* by the amplification of flanking sequences in FH42, FH50 and FH71 strains |
| **OL196** | CTCATCCCGGCAGTTTTC | |
| **OL199** | TGTACGGTTCTGAAGCCAT | *optB* internal primers used to check the absence in FH42, FH50 and FH71 strains |
| **OL200** | GCTGATACTGGATTCCGGAT | |
| **OL203** | ATTGTCGGTGATGTTTCCGG | *fhuE* internal primers used to check the absence in FH43, FH51 and FH72 strains |
| **OL204** | GGTATTCACACGGTCGTTCTG | |
| **OL221** | CATTGCTGGCATGAGCCAG | *optA* internal primers used to check the absence in FH46, FH71, FH72, FH83, FH84 and FH85 strains |
| **OL222** | GAACGTATCCGTGGATGGC | |
| **OL264** | AACGCAAGCTGGAACCGAT | *btuB* amplification by qPCR |
| **OL265** | CTCGCGGATGGTTTCTTCGT | |
| **OL266** | TACAGCGACTGTCCTCGGAT | *1205* amplification by qPCR |
| **OL267** | ACCTCAACCCGCTCTTCGTT | |

| | | |
|---|---|---|
| **OL268** | AGCGCATTGAGGTAGCCAGA | *9625* amplification by qPCR |
| **OL269** | GAGGCAAAAGTGCGCAGGTA | |
| **OL270** | GCAGAACGTTGAGGATCGCT | *optA* amplification by qPCR |
| **OL271** | AACACCGTAGGTCAGGCTGT | |
| **OL272** | GCGTCCTGGAGGATTCTGAT | *asbA* amplification by qPCR [2] |
| **OL273** | ACATGATTGTCCGGCAGGT | |
| **OL274** | TCAGTGCGGAAGTGAATGCT | *optB* amplification by qPCR |
| **OL275** | TTGCGACTATCCGGAGTCAGA | |
| **OL278** | TGGTCCCTGACCGATCAACT | *fhuE* amplification by qPCR |
| **OL279** | GGCATCGTAGCTGCTTTCCT | |
| **OL282** | TGATGACAGCGATGCCAGCA | *rpoD* amplification by qPCR [2] |
| **OL283** | TGTTTGGCCAAGGCCTTCTC | |
| **OL291** | GAAGCTATGAGTACTGCCAAAG | *btuB* internal primers used to check the absence in FH81 and FH84 strains |
| **OL292** | GATATCGTAACGCACGGTCAG | |
| **OL293** | GTCTTGCCCGCGATTTC | Checking Δ*btuB* by the amplification of flanking sequences in FH81 and FH84 strains |
| **OL294** | GTAGTCACTGTAGGCGACC | |
| **OL295** | ACACTTAACGGCTGACATGGGATGAAACCGGTTGGCAAC | Amplification of *btuB*-flanking sequences for pFH77 construct by POE-PCR |
| **OL296** | CCTCCCCCGGTTTAACGGTTTGTTTCACCGCACGCAC | |
| **OL297** | GTGCGTGCGGTGAAACAAACCGTTAAACCGGGGGAGG | |
| **OL298** | GATCTATCAACAGGAGTCCAAGCATCGGTAAACCTGGCTC | |
| **OL299** | GCTCGCTGTATCAGCCTTG | *1205* internal primers used to check the absence in FH82 and FH85 strains |
| **OL300** | GATTGGCCATGAGCGGATG | |
| **OL301** | GCGATGAAGAACGTCCTG | Checking Δ*1205* by the amplification of flanking sequences in FH82 and FH85 strains |
| **OL302** | CACCATCAAGATGGGCG | |
| **OL303** | ACACTTAACGGCTGACATGGTTGATGTGAATTTCACCACTG | Amplification of *1205*-flanking sequences for pFH78 construct by POE-PCR |
| **OL304** | GCAACCGCCATAATCGTTGGGTTGGCGGTCCTTG | |
| **OL305** | CAAGGACCGCCAACCCAACGATTATGGCGGTTGC | |
| **OL306** | GATCTATCAACAGGAGTCCAAGCATCCTGCAATCCGGAAC | |
| **OL307** | GCTCTCTTCATCGCTTGCAAG | *9625* internal primers used to check the absence in FH80 and FH83 strains |
| **OL308** | CCCGGTTCAGGTAATGTTGC | |
| **OL309** | CTCATCGCCTGACGCAAAC | Checking Δ*9625* by the amplification of flanking sequences in FH80 and FH83 strains |
| **OL310** | GATCAGTGGCTCCAGTGTG | |
| **OL311** | ACACTTAACGGCTGACATGGCAACTGGCGACCAATCTC | Amplification of *9625*-flanking sequences for pFH79 construct by POE-PCR |
| **OL312** | GTTTCCTGCCCTGCACACCGGTGTGTCGTTCTATAATTTGCCG | |
| **OL313** | CGGCAAATTATAGAACGACACACCGGTGTGCAGGGCAGGAAAC | |
| **OL314** | GATCTATCAACAGGAGTCCAAGTCAAGGCTTGCCGTAATAAG | |

**Table S4 Amplicon Sequence Variant (ASV) statistics**.

| Experiment | Samples | ASV | | | |
|---|---|---|---|---|---|
| | | # Raw read pairs | # Adapter and primer clipped read pairs | # Combined reads | #No. of reads per sample |
| Polar Front Zone | **a -** Fe-petrobactin | 41 538 | 34 446 | 34 196 | 33 979 |
| | **b -** Fe-petrobactin | 102 988 | 84 033 | 83 686 | 81 989 |
| | **c -** Fe-petrobactin | 25 135 | 20 375 | 20 236 | 20 212 |
| | **a** + Fe-petrobactin | 244 080 | 202 683 | 202 030 | 200 955 |
| | **b** + Fe-petrobactin | 185 181 | 150 189 | 149 719 | 148 143 |
| | **c** + Fe-petrobactin | 69 331 | 54 067 | 53 741 | 51 153 |
| Antarctic Zone | **a -** Fe-petrobactin | 37 801 | 30 190 | 29 912 | 28 534 |
| | **b -** Fe-petrobactin | 70 297 | 57 931 | 57 501 | 51 508 |
| | **c -** Fe-petrobactin | 99 096 | 84 907 | 84 633 | 80 252 |
| | **a** + Fe-petrobactin | 130 720 | 110 115 | 109 555 | 85 593 |
| | **b** + Fe-petrobactin | 108 202 | 90 778 | 90 131 | 72 241 |
| | **c** + Fe-petrobactin | 52 116 | 42 376 | 41 814 | 30 384 |

**Table S6 Sequencing statistics from metagenomes.**
* Co-assembly approach using megahit

| Experiment | Samples | Metagenome | | | | |
|---|---|---|---|---|---|---|
| | | # Read pairs | Avg. contig length (bp)* | # Contigs (≥ 1 000 bp) | Overall alignment rate (%) | No. of genes |
| Polar Front Zone | **a -** Fe-petrobactin | 78 142 245 | 5 994 | 26 263 | 94,58 | 155 490 |
| | **b -** Fe-petrobactin | 84 976 586 | | | 95,86 | |
| | **c -** Fe-petrobactin | 86 551 418 | | | 55,92 | |
| | **a** + Fe-petrobactin | 84 209 812 | | | 98,63 | |
| | **b** + Fe-petrobactin | 88 905 566 | | | 97,87 | |
| | **c** + Fe-petrobactin | 81 889 356 | | | 97,07 | |
| Antarctic Zone | **a -** Fe-petrobactin | 91 958 411 | 7 587 | 41 955 | 94,72 | 311 779 |
| | **b -** Fe-petrobactin | 92 090 723 | | | 92,42 | |
| | **c -** Fe-petrobactin | 96 782 206 | | | 97,72 | |
| | **a** + Fe-petrobactin | 75 909 761 | | | 94,11 | |
| | **b** + Fe-petrobactin | 83 069 294 | | | 90,76 | |
| | **c** + Fe-petrobactin | 95 412 855 | | | 94,05 | |

**Table S7 Sequence comparison of *M. nauticus* TBDTs against the SwissProt database.**
BLASTP 2.16.0+ was used with the BLOSUM45 matrix and the UniProtKB/SwissProt database (573 661 sequences)

| NCBI Gene symbol (Old locus tag) | Protein UniParc ID - Length | Blast best hit: ID - Identity (Query cover) - Length - Organism - Protein name | Best hit with function evidence: ID - Identity (Query cover) - Length - Organism - Protein name |
|---|---|---|---|
| **MARHY_RS16350 (MARHY3432)** | UPI0002E881A7 - 657 AA | **Q2YJB2** - 25.6 % (95 %) - 661 AA - *Brucella abortus* (strain 2308) - Heme transporter BhuA | **Q2YJB2** |
| **MARHY_RS14955 (MARHY3135)** | UPI0002E17DF5 - 676 AA | **A5F9G0** - 38.8 % (99 %) - 652 AA - *Vibrio cholerae* serotype O1 (strain ATCC 39541) - Iron-regulated outer membrane virulence protein | **Q05098** - 25.1 % (96 %) - 746 AA - *Pseudomonas aeruginosa* (strain ATCC 15692 / PAO1) - Ferric enterobactin receptor |
| **MARHY_RS04950 (MARHY1035)** | UPI000256EEEA - 725 AA | **P16869** - 34.1 % (97 %) - 729 AA - *Escherichia coli* (strain K12) - Hydroxamate siderophore receptor FhuE | **P16869** |
| **MARHY_RS01205 (MARHY0250)** | UPI0001DDD475 - 681 AA | **Q66G57** - 21.5 % (85 %) - 624 AA - *Yersinia pseudotuberculosis* serotype I (strain IP32953) - Vitamin B12 transporter BtuB | **P06129** - 19.8 % (85 %) - 614 AA - *Escherichia coli* (strain K12) - Vitamin B12 transporter BtuB |
| **MARHY_RS09625 (MARHY2030)** | UPI000E02E911 - 715 AA | **Q6LLU3** - 23.4 % (66 %) - 606 AA - *Photobacterium profundum* (strain SS9) - Vitamin B12 transporter BtuB | **P06129** - 24.8 % (34 %) - 614 AA - *Escherichia coli* (strain K12) - Vitamin B12 transporter BtuB |
| **MARHY_RS01040 (MARHY0216)** | UPI0001DDD4E6 - 606 AA | **Q6LLU3** - 32,1 % (96 %) - 606 AA - *Photobacterium profundum* (strain SS9) - Vitamin B12 transporter BtuB | **P06129** - 25.4% (95 %) - 614 AA - *Escherichia coli* (strain K12) - Vitamin B12 transporter BtuB |

**Table S8 Global alignments.**

*Matrix BLOSUM 30 ; § Matrix BLOSUM35 ; # Matrix BLOSUM40; Id: Identity ; Sim: Similarity

| | OptA UPI0002E881A7 | OptB UPI0002E17DF5 | FhuE UPI000256EEEA | RS01205 UPI0001DDD475 | RS09625 UPI000E02E911 | BtuB UPI0001DDD4E6 |
|---|---|---|---|---|---|---|
| **OptA UPI0002E881A7** | 100.00 % | | | | | |
| **OptB UPI0002E17DF5** | *Id: 25.2 % Sim: 44.9 % Gaps: 22.0 % | 100.00 % | | | | |
| **FhuE UPI000256EEEA** | *Id: 22.8 % Sim: 40.1 % Gaps: 30.0 % | *Id: 21.3 % Sim: 39.2 % Gaps: 32.0 % | 100.00 % | | | |
| **RS01205 UPI0001DDD475** | *Id: 22.6 % Sim: 41.0 % Gaps: 28.9 % | *Id: 21.5 % Sim: 37.0 % Gaps: 31.6 % | *Id: 19.6 % Sim: 38.6 % Gaps: 30.0 % | 100.00 % | | |
| **RS09625 UPI000E02E911** | *Id: 21.1 % Sim: 36.4 % Gaps: 31.4 % | *Id: 20.7 % Sim: 38.2 % Gaps: 36.5 % | *Id: 19.9 % Sim: 39.7 % Gaps: 30.8 % | *Id: 19.4 % Sim: 35.8 % Gaps: 36.2 % | 100.00 % | |
| **BtuB UPI0001DDD4E6** | *Id: 23.3 % Sim: 40.2 % Gaps: 29.6 % | *Id: 24.6 % Sim: 44.9 % Gaps: 26.1 % | *Id: 22.8 % Sim: 38.6 % Gaps: 34.2 % | *Id: 20.7 % Sim: 40.2 % Gaps: 29.1 % | *Id: 21.7 % Sim: 36.2 % Gaps: 35.5 % | 100.00 % |
| **Q2YJB2** | *Id: 25.6% Sim: 46.2% Gaps: 22.1% | | | | | |
| **A5F9G0** | | #Id: 40.3 % Sim: 57.6 % Gaps: 13.5 % | | | | |
| **Q05098** | | *Id: 27.4 % Sim: 45.6 % Gaps: 20.5 % | | | | |
| **P16869** | | | §Id: 34.6 % Sim: 54.8 % Gaps: 11.7 % | | | |
| **Q66G57** | | | | *Id: 23.1 % Sim: 43.1 % Gaps: 24.6 % | | |
| **P06129** | | | | *Id: 21.0 % Sim: 39.9 % Gaps: 27.6 % | *Id: 21.7 % Sim: 39.8 % Gaps: 27.0 % | *Id: 27.7 % Sim: 50.1 % Gaps: 14.3 % |
| **Q6LLU3** | | | | | *Id: 21.3 % Sim: 38.6 % Gaps: 30.2 % | *Id: 32.6 % Sim: 54.1 % Gaps: 13.8 % |

### Table S9 Summary statistics and taxonomic assignment of the metagenome-assembled genomes (MAGs).

| MAG No. | total length | num. contigs | N50 | GC_content | Completion (%) | Redundancy (%) | t_phylum | t_class | t_order | t_family | t_genus | t_species |
|---|---|---|---|---|---|---|---|---|---|---|---|---|
| **Polar Frontal Zone** | | | | | | | | | | | | |
| **MAG9** | 2128298 | 107 | 27265 | 36.507825954521 | 94,36619718 | 1,408450704 | Pseudomonadota | Alphaproteobacteria | Rhodobacterales | Rhodobacteraceae | Amylibacter | Amylibacter sp905182285 |
| **MAG10** | 3518963 | 487 | 11287 | 34.717325534305 | 92,95774648 | 9,85915493 | Bacteroidota | Bacteroidia | Flavobacteriales | Flavobacteriaceae | Psychroserpens | |
| **MAG13** | 2935249 | 153 | 36116 | 29.110938650567 | 94,36619718 | 7,042253521 | Bacteroidota | Bacteroidia | Flavobacteriales | Flavobacteriaceae | Polaribacter | |
| **MAG15** | 4598165 | 119 | 64168 | 40.491623807827 | 95,77464789 | 5,633802817 | Pseudomonadota | Gammaproteobacteria | Enterobacterales | Alteromonadaceae | Pseudoalteromonas | |
| **MAG17** | 4165794 | 146 | 46193 | 36.235893474360 | 97,18309859 | 0 | Pseudomonadota | Gammaproteobacteria | Enterobacterales | Alteromonadaceae | Colwellia | |
| **MAG20** | 623457 | 120 | 5450 | 33.365086499519 | 57,74647887 | 8,450704225 | Pseudomonadota | Gammaproteobacteria | SAR86 | D2472 | SCGC-AAA076-P13 | |
| **MAG21** | 3191140 | 268 | 24401 | 35.104063919017 | 84,50704225 | 9,85915493 | Bacteroidota | Bacteroidia | Flavobacteriales | Flavobacteriaceae | Ulvibacter | Ulvibacter sp913063405 |
| **MAG22** | 2715987 | 196 | 20908 | 31.483833866776 | 90,14084507 | 1,408450704 | Bacteroidota | Bacteroidia | Flavobacteriales | Flavobacteriaceae | SCGC-AAA160-P02 | SCGC-AAA160-P02 sp000383355 |
| **MAG24** | 3643135 | 73 | 84280 | 64.164473878889 | 100 | 0 | Pseudomonadota | Alphaproteobacteria | Acetobacterales | Acetobacteraceae | Acidocella | Acidocella facilis |
| **MAG26** | 4105585 | 488 | 15838 | 44.118555171190 | 92,95774648 | 16,90140845 | Pseudomonadota | Gammaproteobacteria | Pseudomonadales | Porticoccaceae | HTCC2207 | |
| **MAG28** | 2647390 | 168 | 24373 | 30.113132256671 | 80,28169014 | 1,408450704 | Bacteroidota | Bacteroidia | Flavobacteriales | Flavobacteriaceae | Polaribacter | |
| **MAG29** | 2352556 | 229 | 13723 | 34.940693396940 | 80,28169014 | 2,816901408 | Bacteroidota | Bacteroidia | Flavobacteriales | Flavobacteriaceae | Polaribacter | Polaribacter sp913065125 |
| **MAG44** | 2485400 | 173 | 22182 | 44.829038734466 | 81,69014085 | 5,633802817 | Pseudomonadota | Gammaproteobacteria | Pseudomonadales | Porticoccaceae | HTCC2207 | HTCC2207 sp002313335 |
| **MAG49** | 3210127 | 170 | 29485 | 52.114807369013 | 98,5915493 | 2,816901408 | Pseudomonadota | Gammaproteobacteria | Pseudomonadales | Halieaceae | Halioglobus | Halioglobus sp002862465 |
| **MAG54** | 3759398 | 223 | 25106 | 51.932670556646 | 100 | 0 | Pseudomonadota | Alphaproteobacteria | Rhodobacterales | Rhodobacteraceae | CPC320 | CPC320 sp002336405 |
| **MAG58** | 5175315 | 1078 | 5665 | 30.347277363354 | 97,18309859 | 39,43661972 | Bacteroidota | Bacteroidia | Flavobacteriales | Flavobacteriaceae | Polaribacter | |
| **MAG59** | 1016030 | 125 | 11155 | 34.942835724348 | 76,05633803 | 1,408450704 | Pseudomonadota | Gammaproteobacteria | SAR86 | D2472 | SCGC-AAA076-P13 | |
| **MAG67** | 4523165 | 1117 | 4173 | 44.344402714767 | 87,32394366 | 26,76056338 | Pseudomonadota | Alphaproteobacteria | Rhodobacterales | Rhodobacteraceae | Planktomarina | |
| **MAG69** | 4092574 | 505 | 11819 | 36.989505000134 | 91,54929577 | 14,08450704 | Pseudomonadota | Gammaproteobacteria | Enterobacterales | Alteromonadaceae | Colwellia | |
| **MAG70** | 2977552 | 16 | 295225 | 43.906768585777 | 98,5915493 | 0 | Pseudomonadota | Gammaproteobacteria | Nitrosococcales | Methylophagaceae | Methylophaga | |
| **MAG72** | 2530125 | 459 | 6814 | 50.986461953148 | 95,77464789 | 8,450704225 | Pseudomonadota | Alphaproteobacteria | Rhodobacterales | Rhodobacteraceae | Ascidiaceihabitans | |
| **MAG76** | 2817500 | 284 | 15623 | 42.485997140372 | 100 | 2,816901408 | Pseudomonadota | Gammaproteobacteria | Pseudomonadales | | | |
| **MAG81** | 4702365 | 138 | 55539 | 36.374265894824 | 85,91549296 | 1,408450704 | Pseudomonadota | Gammaproteobacteria | Enterobacterales | Alteromonadaceae | Colwellia | |
| **MAG82** | 3895059 | 248 | 27974 | 37.097632917176 | 100 | 4,225352113 | Pseudomonadota | Gammaproteobacteria | Enterobacterales | Alteromonadaceae | | |
| **MAG86** | 2672666 | 474 | 6834 | 44.005207422608 | 97,18309859 | 0 | Pseudomonadota | Gammaproteobacteria | Pseudomonadales | Moraxellaceae | Psychrobacter | |
| **Antarctic Zone** | | | | | | | | | | | | |
| **MAG3** | 2795192 | 69 | 106181 | 44.631666759429 | 98,5915493 | 7,042253521 | Pseudomonadota | Gammaproteobacteria | Pseudomonadales | Nitrincolaceae | ASP10-02a | ASP10-02a sp002312935 |
| **MAG5** | 3116984 | 153 | 41837 | 30.503658963260 | 88,73239437 | 7,042253521 | Bacteroidota | Bacteroidia | Flavobacteriales | Flavobacteriaceae | | |
| **MAG10** | 3901581 | 320 | 17173 | 54.551040728039 | 98,5915493 | 4,225352113 | Pseudomonadota | Alphaproteobacteria | Rhodobacterales | Rhodobacteraceae | Ascidiaceihabitans | Ascidiaceihabitans sp905182355 |
| **MAG16** | 4444187 | 153 | 61509 | 38.473065691058 | 95,77464789 | 2,816901408 | Pseudomonadota | Gammaproteobacteria | Enterobacterales | Psychromonadaceae | Psychromonas | Psychromonas sp002835995 |
| **MAG22** | 3142745 | 287 | 21580 | 44.852602340508 | 98,5915493 | 1,408450704 | Pseudomonadota | Gammaproteobacteria | Pseudomonadales | Porticoccaceae | HTCC2207 | |
| **MAG33** | 2310430 | 52 | 69612 | 30.987551833824 | 98,5915493 | 0 | Bacteroidota | Bacteroidia | Flavobacteriales | Flavobacteriaceae | GCA-002733185 | GCA-002733185 sp913061205 |
| **MAG36** | 2378227 | 142 | 53243 | 35.055000898951 | 98,5915493 | 1,408450704 | Bacteroidota | Bacteroidia | Flavobacteriales | Flavobacteriaceae | Patiriisocius | Patiriisocius sp003450155 |
| **MAG38** | 5311933 | 415 | 23617 | 57.624418548178 | 87,32394366 | 8,450704225 | Pseudomonadota | Alphaproteobacteria | Rhodobacterales | Rhodobacteraceae | Sulfitobacter | Sulfitobacter pontiacus |
| **MAG40** | 2245020 | 136 | 26069 | 31.910200030502 | 97,18309859 | 0 | Bacteroidota | Bacteroidia | Flavobacteriales | Flavobacteriaceae | SCGC-AAA160-P02 | SCGC-AAA160-P02 sp000383355 |
| **MAG48** | 3087556 | 280 | 14103 | 47.689939502171 | 84,50704225 | 0 | Pseudomonadota | Gammaproteobacteria | Pseudomonadales | HTCC2089 | UBA4582 | UBA4582 sp905182025 |
| **MAG54** | 6117841 | 35 | 327886 | 39.575983190399 | 100 | 2,816901408 | Pseudomonadota | Gammaproteobacteria | Enterobacterales | Alteromonadaceae | Paraglaciecola | |
| **MAG57** | 2185443 | 91 | 24021 | 38.118755216586 | 63,38028169 | 4,225352113 | Bacteroidota | Bacteroidia | Flavobacteriales | Flavobacteriaceae | MAG-121220-bin8 | MAG-121220-bin8 sp905182765 |
| **MAG64** | 4660140 | 235 | 28503 | 40.720409821173 | 94,36619718 | 25,35211268 | Pseudomonadota | Gammaproteobacteria | Enterobacterales | Alteromonadaceae | Pseudoalteromonas | |
| **MAG66** | 2424972 | 705 | 3501 | 36.081725223747 | 70,42253521 | 5,633802817 | Bacteroidota | Bacteroidia | Flavobacteriales | Flavobacteriaceae | Maribacter | |
| **MAG73** | 2501198 | 50 | 89322 | 48.468999745971 | 84,50704225 | 1,408450704 | Pseudomonadota | Gammaproteobacteria | Pseudomonadales | Porticoccaceae | HTCC2207 | HTCC2207 sp905182855 |
| **MAG76** | 4583881 | 1085 | 4510 | 62.229660827550 | 87,32394366 | 22,53521127 | Pseudomonadota | Alphaproteobacteria | Rhodobacterales | Rhodobacteraceae | Sulfitobacter | |
| **MAG78** | 4942799 | 63 | 124461 | 38.353615091948 | 91,54929577 | 0 | Pseudomonadota | Gammaproteobacteria | Pseudomonadales | Nitrincolaceae | Rs1 | |
| **MAG79** | 3187265 | 75 | 128343 | 54.898585654254 | 73,23943662 | 2,816901408 | Pseudomonadota | Alphaproteobacteria | Rhodobacterales | Rhodobacteraceae | Yoonia | Yoonia sp002378665 |
| **MAG81_2** | 2585287 | 51 | 101716 | 31.481938217233 | 98,5915493 | 2,816901408 | Bacteroidota | Bacteroidia | Flavobacteriales | Flavobacteriaceae | Polaribacter | Polaribacter sp913061165 |
| **MAG82** | 3061822 | 439 | 9656 | 30.910639755796 | 95,77464789 | 11,26760563 | Bacteroidota | Bacteroidia | Flavobacteriales | Flavobacteriaceae | Polaribacter | |
| **MAG92** | 3935753 | 459 | 13880 | 43.808602302396 | 95,77464789 | 8,450704225 | Pseudomonadota | Gammaproteobacteria | Thiotrichales | Thiotrichaceae | Leucothrix | Leucothrix arctica |
| **MAG95** | 2405559 | 267 | 11949 | 30.391294563793 | 85,91549296 | 14,08450704 | Bacteroidota | Bacteroidia | Flavobacteriales | Flavobacteriaceae | SGZJ01 | SGZJ01 sp913009535 |
| **MAG102** | 4487725 | 379 | 17008 | 53.926517185889 | 100 | 12,67605634 | Pseudomonadota | Alphaproteobacteria | Rhodobacterales | Rhodobacteraceae | Sulfitobacter | |
| **MAG107** | 3442223 | 93 | 62204 | 40.454222487535 | 98,5915493 | 7,042253521 | Pseudomonadota | Gammaproteobacteria | Pseudomonadales | Marinomonadaceae | Marinomonas | Marinomonas colpomeniae |

#### Table S11 TBDT used in Figure 7.

| Organism Name | UniParc ID - Mature protein position | UniProt-RefSeq-ID | Name | Siderophores transported | Siderophores produced | Reference |
|---|---|---|---|---|---|---|
| *Alteromonas macleodii* ATCC 27126 | UPI0001AEBB3D 21-681 AA | WP_014951117 | | Unknown | Petrobactin | [22] |
| *Aminobacter aminovorans* KCTC_2477 | UPI00078D4168 43-768 AA | A0AAC8YJ23 | | Unknown | Unknown | |
| *Aureimonas sp.* AU20 | UPI00071EFED4 29-755 AA | A0A0S2EV74 | | Unknown | | |
| *Azospirillum ramasamyi M2T2B2* | UPI000DA33E59 18-739 AA | A0A2U9SFF3 | | Unknown | | |
| *Ensifer adhaerens Casida A-T305* | UPI0004A95805 47-768 AA | A0ABY8HN84 | | Unknown | Unknown | |
| *Flavobacterium collinsii TRV642* | UPI0021DFF913 22-823 | A0A9W4TK89 | BtuB | Vitamin B12 | Unknown | |
| ***Escherichia coli* K12** | | | | | Enterobactin | [29] |
| *Escherichia coli* K12 | UPI000016EE5E 26-663 AA | P17315 | CirA | Catecholates | | [30] |
| *Escherichia coli* K12 | UPI000012A647 23-746 AA | P05825 | FepA | Enterobactin | | [31] |
| *Escherichia coli* K12 | UPI000013A36D 32-760 AA | P75780 | Fui | Catecholates | | [32] |
| *Leisingera methylohalidivorans* MB2 | UPI0003D69F8B 29-661 AA | V9VTN1 | | Unknown | Unknown | |
| ***Marinobacter nauticus* SP17** | | | | | Petrobacin, petrobactin monosulfonate and disulfonate | [33] |
| *Marinobacter nauticus* SP17 | UPI000256EEEA 29-725 AA | WP_014420810 | FhuE | Unknown | | |
| *Marinobacter nauticus* SP17 | UPI0002E17DF5 25-676AA | A0A368Y2X9 | OptB | Petrobactin | | This work |
| *Marinobacter nauticus* SP17 | UPI0002E881A7 30-657 AA | WP_041655120 | OptA | petrobactin | | This work |
| *Marinobacter shengliensis D49* | UPI00257419E0 31-657 AA | WP_286252133 | | Unknown | | |
| *Mesorhizobium sp.* AA22 | UPI0007ED2DF4 26-695 AA | A0A8T8D0F2 | | Unknown | | |
| *Methylorubrum extorquens* AM1 | UPI0001A7A532 22-839 AA | C5B1I1 | | Unknown | Methylolanthanin | [34] |
| *Microbulbifer sp.* A4B17 | UPI0018640994 27-694 AA | WP_192879356 | | Unknown | | |
| ***Nitrobacter winogradskyi* Nb-255** | | | | | Unknown | |
| *Nitrobacter winogradskyi* Nb-255 | UPI00005B379A 27-773 AA | Q3SQ93 | | Unknown | | |
| *Nitrobacter winogradskyi* Nb-255 | UPI00005B379C 32-752 AA | Q3SQ91 | | Unknown | | |
| *Phaeobacter gallaeciensis* DSM 26640 | UPI0003D6D6D2 33-664 AA | A0AAD0EF25 | | Unknown | Unknown | |
| *Phaeobacter inhibens* 2.10 | UPI000160CCC6 33-664 AA | WP_014889464 | | Unknown | Roseobactin | [35] |
| *Photobacterium swingsii CECT7576* | UPI000D1872DC 27-644 AA | A0A2T3NXI7 | | Unknown | | |
| *Pseudomonas aeruginosa LESB58* | UPI0001E9E184 36-766 AA | A0AAQ3QYH1 | PiuD | Catechol siderophore drug conjugates | | [36] |
| ***Pseudomonas aeruginosa* PAO1** | | | | | Pyoverdine, pyochelin | [37] |
| *Pseudomonas aeruginosa* PAO1 | UPI00000D4235 36-753 AA | G3XCY8 | PiuA | Catecholamines | | [38] |
| *Pseudomonas* | UPI00000C51C4 | Q9I527 | PirA | Enterobactin | | [39] |

| | | | | | | |
|---|---|---|---|---|---|---|
| *aeruginosa* PAO1 | 29-742 AA | | | Catecholates | | |
| *Pseudomonas aeruginosa* PAO1 | UPI000013174B 26-746 AA | Q05098 | PfeA | Enterobactin | | [40] |
| *Pseudomonas aeruginosa* PAO1 | UPI00000C56EB 48-820 AA | Q9I116 | FoxA | Ferrioxamine B Nocardamine | | [41] |
| *Pseudomonas aeruginosa PAO1* | UPI00000C5C4D 23-802 AA | Q9HWL3 | FpvB | Pyoverdine Hydroxamates | | [42] |
| *Pseudomonas aeruginosa PAO1* | UPI000012ABCB 44-815 AA | P48632 | FpvA | Pyoverdine | | [43] |
| *Rhodopseudomonas palustris* DX-1 | UPI0001C26DF1 32-754 AA | E6VNP2 | | Unknown | Unknown | |
| ***Rhodopseudomonas palustris* CGA009** | | | | | Rhodopetrobactin A and B | [44] |
| *Rhodopseudomonas palustris* CGA009 | UPI000022705A 28–771 AA | Q6N782 | | Unknown | | |
| *Rhodopseudomonas palustris* CGA009 | UPI000022705C 29–743 AA | Q6N780 | | Unknown | | |
| *Rhodopseudomonas palustris* CGA009 | UPI000022706C 39–797 AA | Q6N764 | | Unknown | | |
| *Sinorhizobium fredii* HH103 | UPI000241767E 47-767 AA | G9A5Y5 | | Unknown | Unknown | |
| | | | **TBDT sequences from MAG** | | | |
| *Pseudoalteromonas sp.* | SIDEMAR MAG 15 | | OptB-like | Unknown | Unknown | This work |
| *Pseudoalteromonas sp.* | SIDEMAR MAG 64 | | OptB-like | Unknown | Unknown | This work |
| *Halopseudomonas pachastrellae* | AF82bin23_orig_MAG301 27-688 | | OptB-like | Unknown | Unknown | [45] |
| *Pseudoalteromonas atlantica* | AF102bin16_orig_MAG64 32-690 | | OptB-like | Unknown | Unknown | [45] |
| *Pseudoalteromonas gelatinilytica* | AF83bin6_strict_MAG312 26-658 | | OptB-like | Unknown | Unknown | [45] |
| *Halomonas titanicae* | AF132bin10_strict_MAG245 25-654 | | OptB-like | Unknown | Unknown | [45] |
| *Halopseudomonas oceani* | AF82bin23_orig_MAG301_2 25-745 | | OptB-like | Unknown | Unknown | [45] |
| *Halomonas titanicae* | AF132bin10_strict_MAG245_2 25-710 | | OptB-like | Unknown | Unknown | [45] |
| *Sphingobium yanoikuyae* | AF95bin8_orig_MAG345 25-654 | | OptB-like | Unknown | Unknown | [45] |

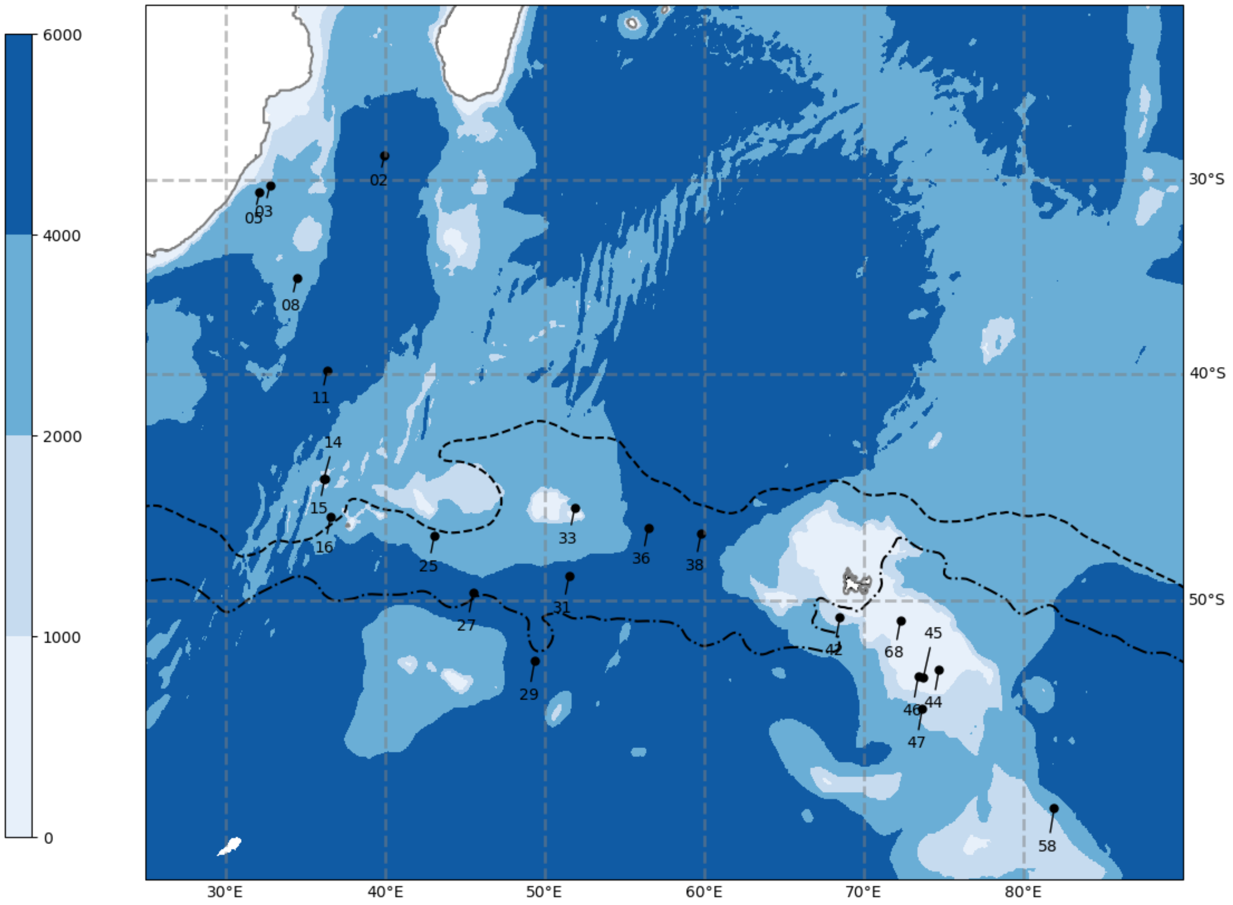


**Figure S1 Map of stations from the SWINGS cruise in the Indian Sector of the Southern Ocean.** Shown are stations where water was collected for the incubation experiments (Station 27 and Station 47) and for metagenomes [45]. Station 27 is located in the Polar Frontal Zone (PFZ; 45.525E, 49.714S) and Station 47 in the Antarctic Zone (AAZ; 73.664E, 54.169S). Concentrations of DOC were 49.6 ± 1 µM 45.5 ± 0.1 µM in surface waters at Station 27 and 47, respectively and of dissolved Fe 0.49 ± 0.08 nM and 0.21 ± 0.0 nM at Station 27 and 47, respectively. Colour shading represents bathymetry and the grey line contours South Africa (25°–35° E) and Madagascar (45°–50° E). The subantarctic front and polar front are drawn according to Park et al. [46].

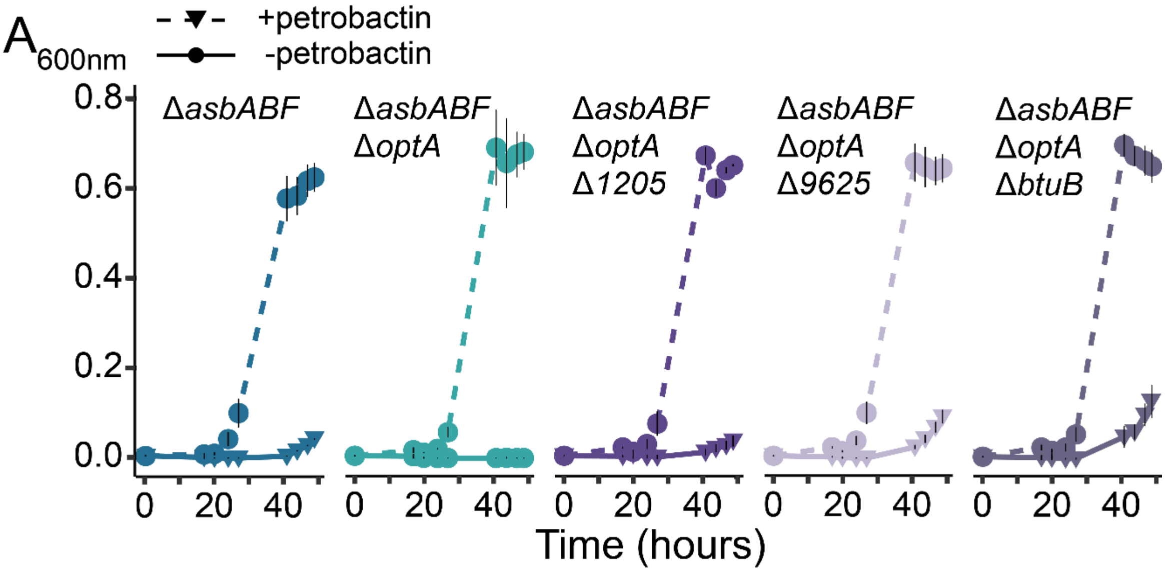


**Figure S2 TBDT deletion mutants supplementation with petrobactin.** Growth curves of TBDT deletion mutant in the absence or the presence of petrobactin at 10 µM. Growth is displayed as $A_{600nm}$ measurements in function of time in hours and correspond to the mean (± SD) of three independent experiments. Δ*asbABF* (FH35), Δ*asbABF* Δ*optA* (FH46), Δ*asbABF* Δ*optA* Δ*9625* (FH83), Δ*asbABF* Δ*optA* Δ*btuB* (FH84), Δ*asbABF* Δ*optA* Δ*1205* (FH85)

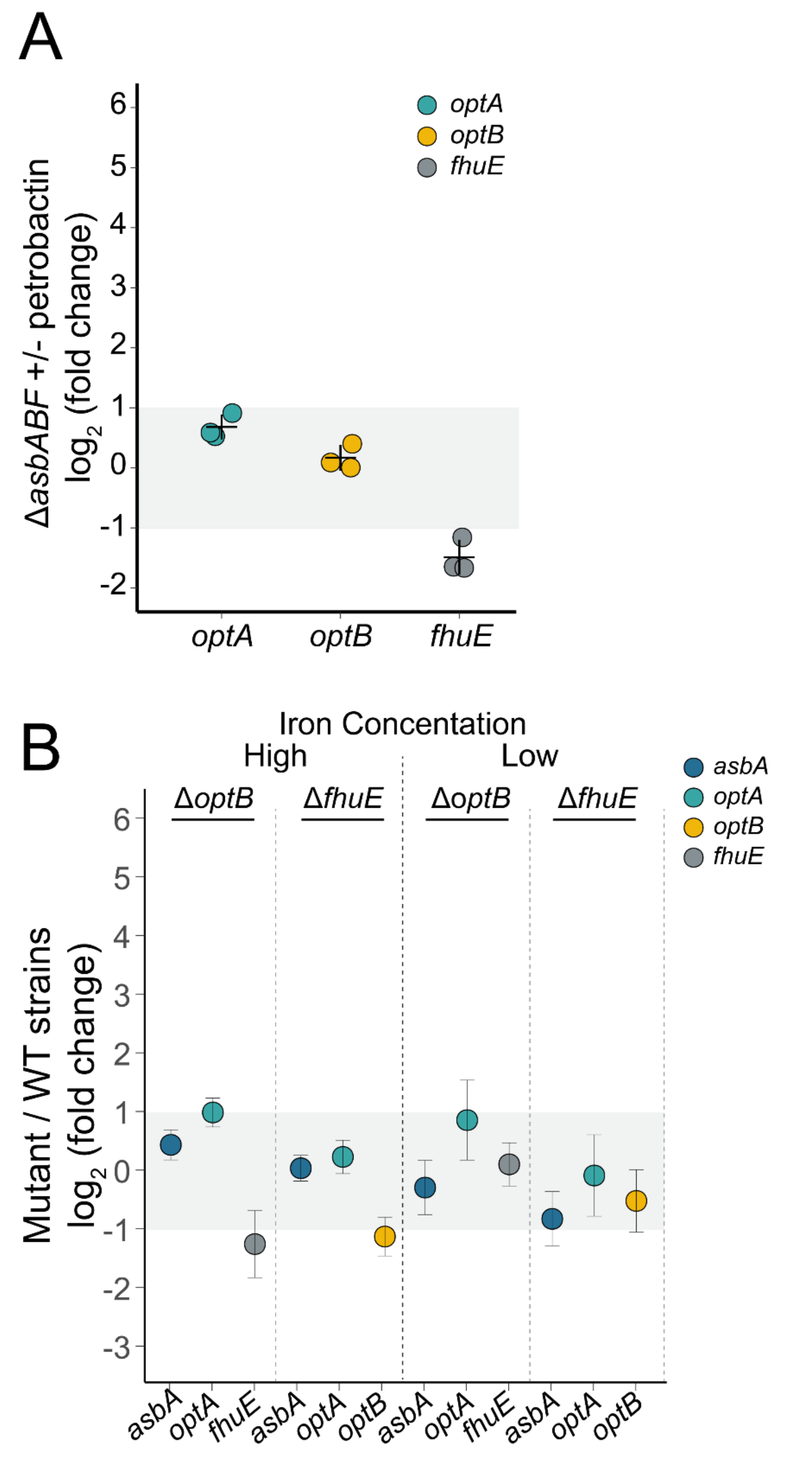


**Figure S3 Relative expression levels of TBDTs and petrobactin biosynthesis genes measured by RT-qPCR.** Results are shown as dot plots based on data from at least three biological replicates (plain dots). RT-qPCR reactions for each sample were carried out in triplicate. Relative fold changes in gene expression were calculated according to the $2^{\Delta\Delta Ct}$ method. None of $log_2$ fold change values not significantly different from 1, 0 or -1 (p-value $< 0.05$, one-sample t-test). **(A)** Expression ratios of *optA*, *optB*, *fhuE* in presence versus absence of 10 µM petrobactin in Δ*asbABF* background, under 1 µM iron concentration and 200 µM EDTA. Error bar represent mean ± standard deviation (SD). **(B)** Expression ratios of *asbA*, *optA*, *optB* and *fhuE* in Δ*optB or* Δ*fhuE* versus WT backgrounds under low and high iron concentration SD was estimated from the distribution of fold changes obtained by 2 000 bootstrap resampling iterations, and error bars represent mean ± SD.

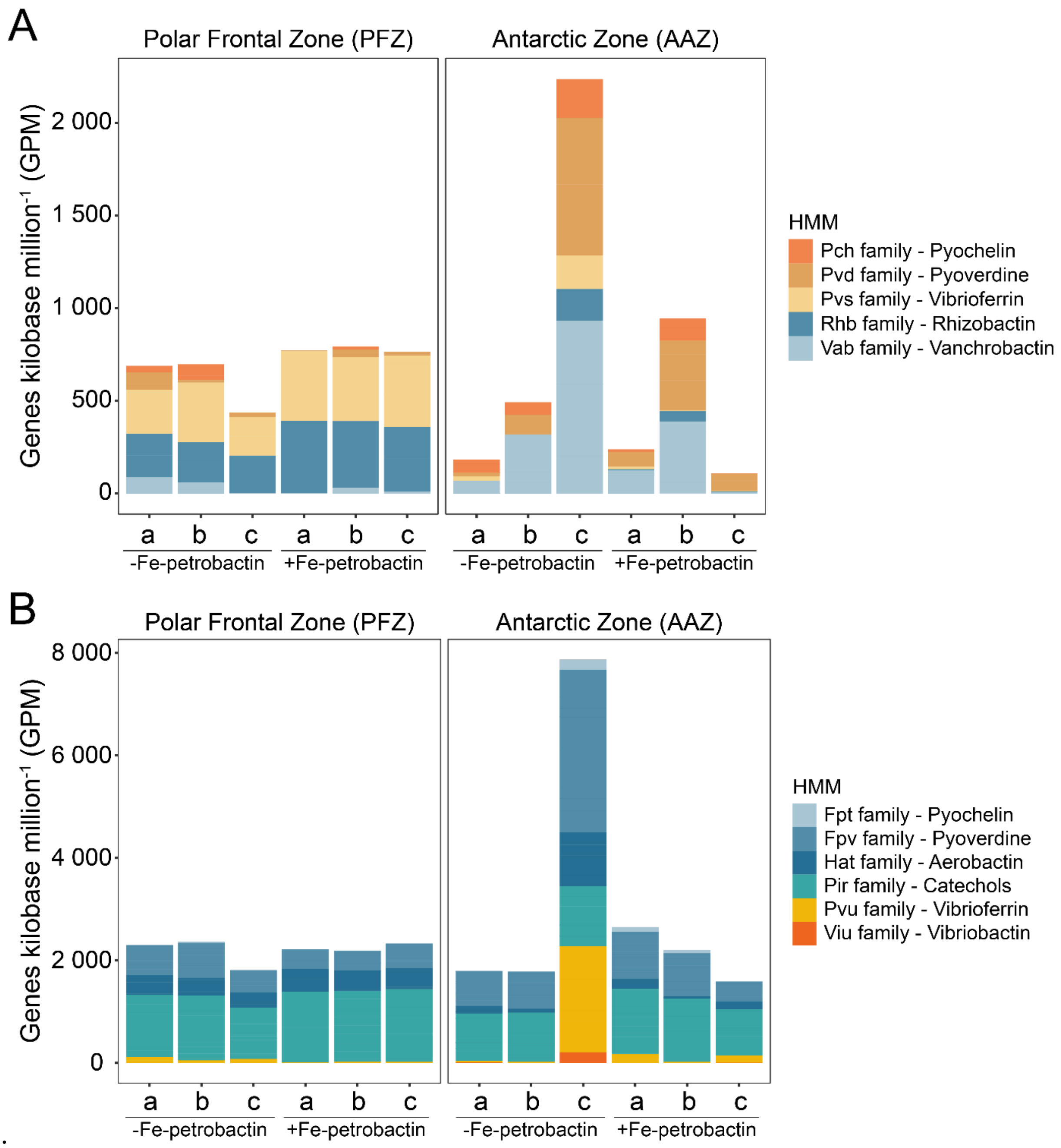


.

**Figure S4 GPMs of siderophore biosynthesis and transport at the end of the incubation experiments.** Genes per kilobase million (GPM) for the subgenes detected in the metagenomes are shown. (**A**) Siderophore synthesis. Pch: *pchBCGHI*; Pvd: *pvdEQRT*; Pvs: *pvsACDE*; Rhb: *rhbABDE*; Vab: *vabABCEFGH*. (**B**) Siderophore transport. Fpt: *fptCX*; Fpv: *fpvCDEFGJK;* Hat: *hatD*; Pir: *pirA*; Pvu: *pvuBCDE;* Viu: *viuB*.

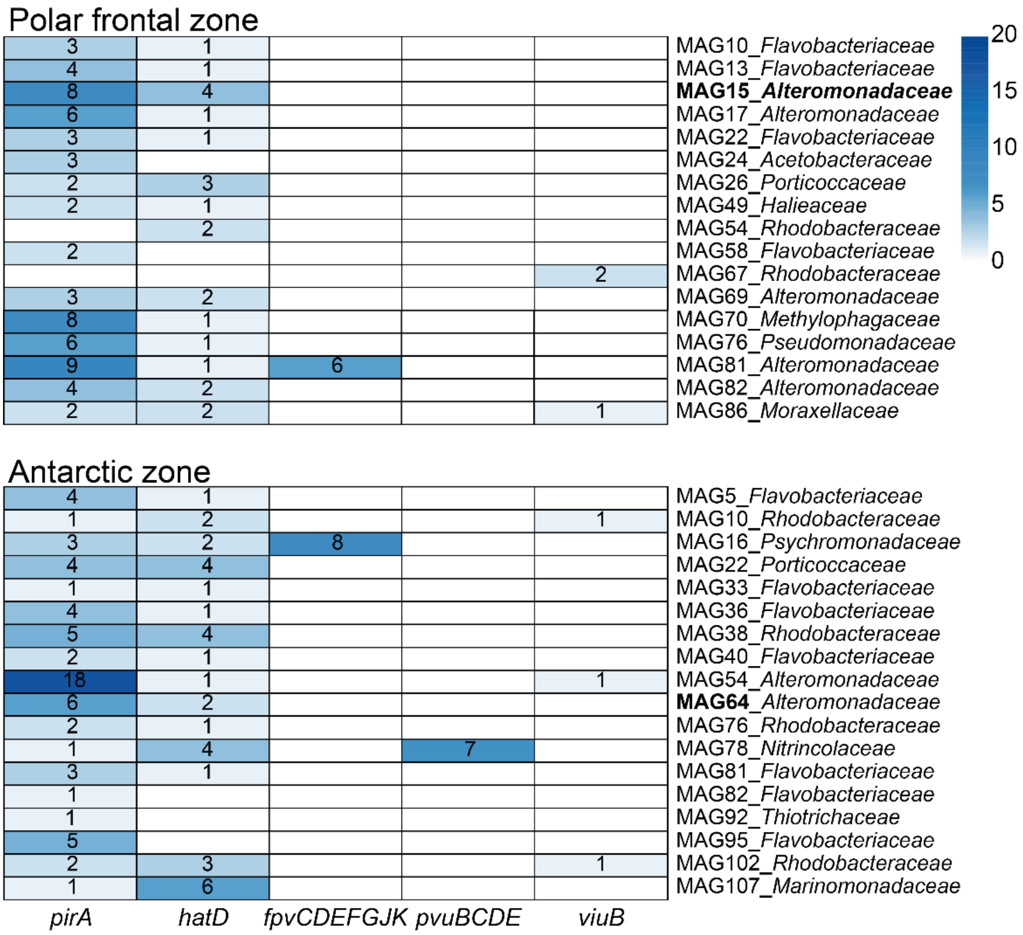


**Figure S5 Siderophore transport genes count.** Heatmap displaying the gene counts of siderophore transport genes annotated as different subgenes that were detected in MAGs reconstructed from metagenomes from the two experiments: Polar frontal zone and Antarctic zone. Encoded proteins predicted functions (and localisation) are as followed: PirA - TBDT (outer membrane); HatD - ABC transporter (inner membrane); FpvCF - substrate binding proteins (periplasmic); FpvDE - ABC transporter (inner membrane); FpvGJK - inner membrane transporter; PvuBCDE - ABC transporter (inner membrane); ViuB – (cytosolic protein)

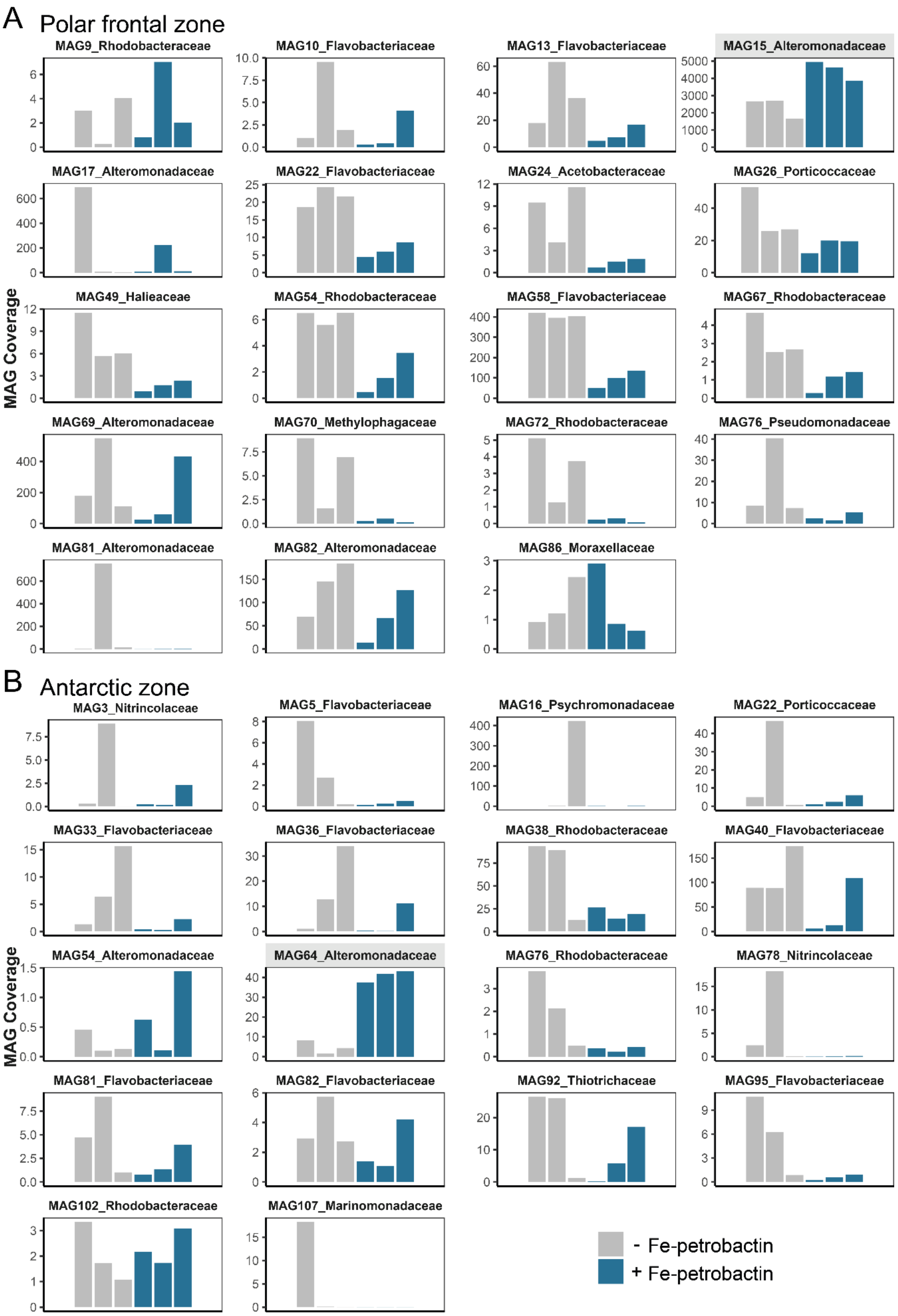


**Figure S6 Coverage of MAGs obtained from incubation experiments.** Bar graphs represent the coverage of MAG for each replicate of the control (- Fe-petrobactin; grey) and amended treatment (+ Fe-petrobactin; dark blue) in the Polar frontal zone (**A**) and Antarctic zone (**B**) experiments.

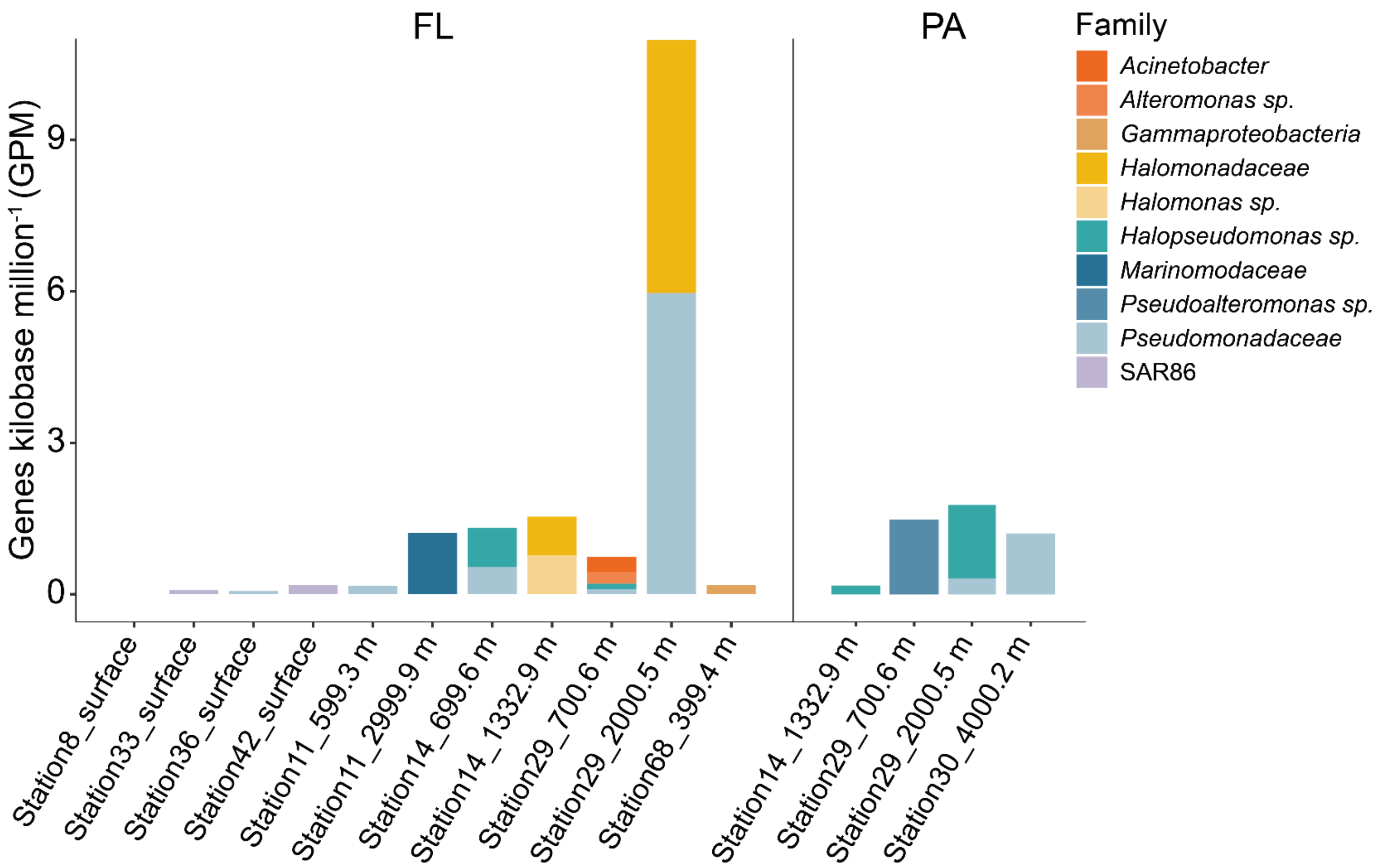


**Figure S7 Relative abundance of *optB*-like genes in metagenomes.** Metagenomes of seawater were collected across the SWINGS cruise transect. *optB*-like genes were searched for in a total of 23 metagenomes in the free-living (FL, < 0.8 µm size fraction) fraction and 19 metagenomes in the particle-attached (PA, > 0.8 µm size fraction) fraction [45].
